# Development of an Enclosed Microbial Co-culture Device and GC×GC-TOFMS Workflow to Investigate Volatile Organic Compound Exchange

**DOI:** 10.64898/2026.11.13.766554

**Authors:** Ting Wang, Wenjing Ma, Jamison M. Whitten, Jodie C. Tokihiro, Hailey Chadrow, Jaimie Choi, Keila Y. Uchimura, Ulri N. Lee, Breanne N. Steffan, Nancy P. Keller, Erwin Berthier, Robert E. Synovec, Ashleigh B. Theberge, Lauren J. Rajakovich

## Abstract

Microbial volatile organic compounds (mVOCs) can mediate interactions between microorganisms without direct physical contact. However, co-culture systems to examine these interactions are limited, and volatilomic data interpretation is challenging. Here, we developed enclosed microbial co-culture platforms compatible with diverse microbial systems, headspace solid-phase microextraction (HS-SPME), and comprehensive two-dimensional gas chromatography time-of-flight mass spectrometry (GC×GC-TOFMS) for mVOC sampling and characterization. In a double-layer co-culture device, diffusion of exogenously supplemented *Aspergillus fumigatus* mVOCs, 1-octen-3-ol and isopentyl alcohol, inhibited *Pseudomonas aeruginosa* growth and biofilm formation. The device was then used to examine mVOC-mediated interkingdom interactions between the bacterium *P. aeruginosa* and the spore-forming fungus *A. fumigatus*. GC×GC-TOFMS was used to profile mVOCs in four sample classes: media, each individual monoculture, and co-culture. Tile-based F-ratio analysis identified 198 analyte hits that differed across at least two of the four sample classes. Subsequent ANOVA identified 104 of these analytes as statistically significant (p < 0.01). To systematically classify their abundance patterns, pairwise comparisons among the four sample classes were encoded using a six-digit ternary code representing the direction and statistical significance of each difference. Of the 104 significant analytes, 88 were assigned to 15 recurring ternary-code groups, capturing distinct patterns associated with microbial production, suppression in co-culture, and differential consumption of media-derived compounds. Closely related ternary-code groups were further consolidated into five major Categories, providing biologically interpretable mVOC profiles. Overall, the integrated co-culture platform and analytical approach provide a versatile framework for investigating microbial volatile interactions and interpreting complex multidimensional volatilomic datasets.

**Table of Contents Graphic:** 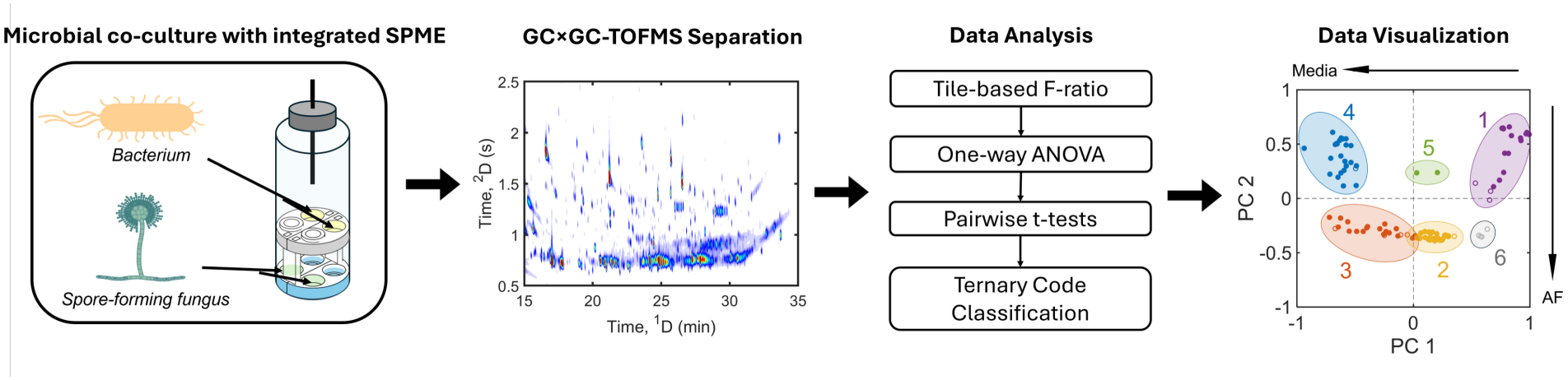

## INTRODUCTION

Microorganisms such as bacteria and fungi produce chemical signals as a primary form of inter- and intra-kingdom communication. These metabolites can be exchanged by direct cell-cell contact, at short distances via diffusible chemicals, or over long distances and physical barriers via volatile signals.^1–4^ Microbial volatile organic compounds (mVOCs) are a diverse group of small molecules and metabolites (<300 Da) that can diffuse in the air, enabling communication with neighbors without direct physical contact.^2,5^ Volatile-mediated communication occurs in diverse microbial environments, including the gut, lungs, soil and microbial mats where microorganisms coexist within complex communities, driving synergistic or antagonistic outcomes. To study mVOC-mediated communication, two major challenges need to be addressed: 1) a lack of controlled experimental systems for isolating and examining volatile interactions in co-cultures and 2) a lack of robust analytical approaches for efficient sampling and systematic characterization of complex microbial volatilomic datasets.

Several co-culture platforms have been developed for spatially separated microbial co-cultivation, such as the widely used split Petri dishes (BI-Petri dish) and two-Petri-dish (plate-in-plate) systems.^6–8^ However, these systems require a secondary container to minimize volatile loss, limiting their use in volatile collection and sampling. A second major limitation of these systems is their incompatibility with diverse microorganisms, particularly spore-forming microbes. Airborne spores can diffuse through the shared headspace, bypassing the intended spatial barriers and potentially introducing physical interactions that interfere with mVOC characterization or make it challenging to understand contributions from volatile signaling versus physical interactions. A double-flask chamber system, in which each microbial culture is enclosed in a glass flask and connected with glass tubing, has also been developed to assess volatile communication.^9^ Despite providing improved volatile collection and physical separation, it has limited scalability for biological replicates due to large footprints and requires additional handling and labor for volatile sampling. Therefore, there is a need for a simple and versatile co-culture system that supports diverse microbes, maintains physical separation, isolates headspace mVOC exchange, and facilitates efficient mVOC sampling for easy integration with workflows based upon gas chromatography-mass spectrometry (GC-MS) and related instrumental methods.

Comprehensive two-dimensional (2D) gas chromatography coupled with time-of-flight mass spectrometry (GC×GC-TOFMS) is a powerful approach for characterizing complex VOC profiles in biological systems. Its enhanced chromatographic separation and mass spectrum acquisition enable broad, untargeted characterization of chemically diverse samples.^10–14^ Accordingly, GC×GC-TOFMS has been applied to investigate metabolic and VOC changes associated with disease and infection,^13–16^ diet and breath profiles,^9,10^ microbial metabolism and volatilomes,^17–31^ and other biological systems.^24–26^ The ability to simultaneously resolve and detect a large number of compounds is particularly valuable for mVOC analysis, where multiple compounds can be produced, consumed, or altered across biological conditions. However, this analytical advantage also produces highly complex datasets containing hundreds to thousands of features distributed across the 2D chromatographic space, making comprehensive manual interpretation impractical. Chemometric approaches are therefore essential for efficiently extracting chemically informative features and identifying patterns associated with different sample classes.

Chemometric analysis of chromatographic data can be broadly divided into targeted and nontargeted approaches. Targeted analysis focuses on predefined chemical compounds or features, whereas nontargeted analysis seeks to identify chemical features associated with differences or similarities among samples without requiring their identities to be specified *a priori*.^27–31^ Nontargeted approaches can be further categorized as supervised or unsupervised, depending on whether sample-class information is incorporated into the analysis. Among supervised nontargeted methods, tile-based Fisher-ratio (F-ratio) analysis provides an effective approach for identifying class-discriminating analytes in GC×GC datasets.^30–36^ In this approach, the F-ratio is calculated as the ratio of between-class variance to within-class variance for a chromatographic tile on a per mass-channel (*m/z*) basis. The resulting analyte hits are ranked according to their F-ratios, allowing features most likely to distinguish sample classes to be prioritized for further investigation. Selected features can further be evaluated using multivariate approaches such as principal component analysis (PCA) to visualize relationships among samples and analyte response patterns.^36,37^ However, when multiple sample classes are considered, analytes can exhibit diverse and overlapping response patterns not readily captured by conventional visualization alone, motivating additional approaches for systematic classification of analyte hits.

In this study, we developed two microbial volatile co-culture platforms enclosed in 20 mL GC-MS glass vials, which can accommodate both non-spore-forming and spore-forming microbes. Both devices are compatible with headspace (HS) solid-phase microextraction (SPME) followed by GC×GC-TOFMS, enabling efficient sampling and characterization of mVOC profiles across diverse co-culture conditions. We validated the capability of our double-layer device to isolate and encapsulate volatile-mediated interactions. We further applied the co-culture platform to study mVOC communication and collect microbial volatilomes employing the opportunistic fungus *Aspergillus fumigatus* and pathogenic bacterium *Pseudomonas aeruginosa* as a model co-culture system. We then acquired HS-SPME-GC×GC-TOFMS data to characterize mVOC profiles in the co-culture system. The dataset comprised four sample classes: *P. aeruginosa* (PA), *A. fumigatus* (AF), their co-culture (CC), and a media control (M), with six biological replicates per class. Tile-based F-ratio analysis was used to identify analytes exhibiting statistically significant differences among the four sample classes. We then developed a probability-based ternary-code approach to systematically classify the diverse response patterns of these analytes. PCA and ternary-code patterns were used to organize class-discriminating analytes according to shared response patterns and facilitated interpretation of VOCs associated with individual microorganisms, co-culture, and media conditions. The novel co-culture devices and systematic analytical approach extend our ability to understand mVOC-mediated inter-species interactions.

## EXPERIMENTAL SECTION

### mVOC co-culture device design and fabrication

Computer-aided designs (CAD) and computer-aided machining (CAM) G-code (.simpl) files for the single-layer and double-layer co-culture device were created in Autodesk Fusion (Autodesk, San Francisco, CA). Devices were milled using a Datron Neo computer numerical control (CNC) mill (Datron Dynamics, Milford, New Hampshire) in 4 mm polystyrene sheets (#ST31-SH-000200; Goodfellow Corporation, Pittsburgh, PA). The culture layers of the double-layer device were milled on both sides of the polystyrene sheet using 1/32” (TR-2-0312-BN) and 1/64” (TR-2-0150-BN)

(Performance Micro Tool; Janesville, WI) diameter endmills. After fabrication, the devices were ultrasonicated in 70% ethanol for 30 min and dried with compressed air. The 1 μm polytetrafluoroethylene (PTFE) membranes (The Lab Depot, Dawsonville, GA) were trimmed using a cutting plotter (CE6000-40; Graphtec Inc., Irvine, CA) based on a design generated in Adobe Illustrator 2025 (Figure 1B). Prior to assembly, all components of the device were sterilized by UV irradiation for 20 min on each side. More details on device dimensions can be found in the Supporting Information (Figures S1 and S2); CAD files for the polystyrene devices and the PDF (editable in Adobe Illustrator) design file used for the plotter cuter to cut the PTFE membranes are included in the electronic supplemental files.

**Figure 1.**
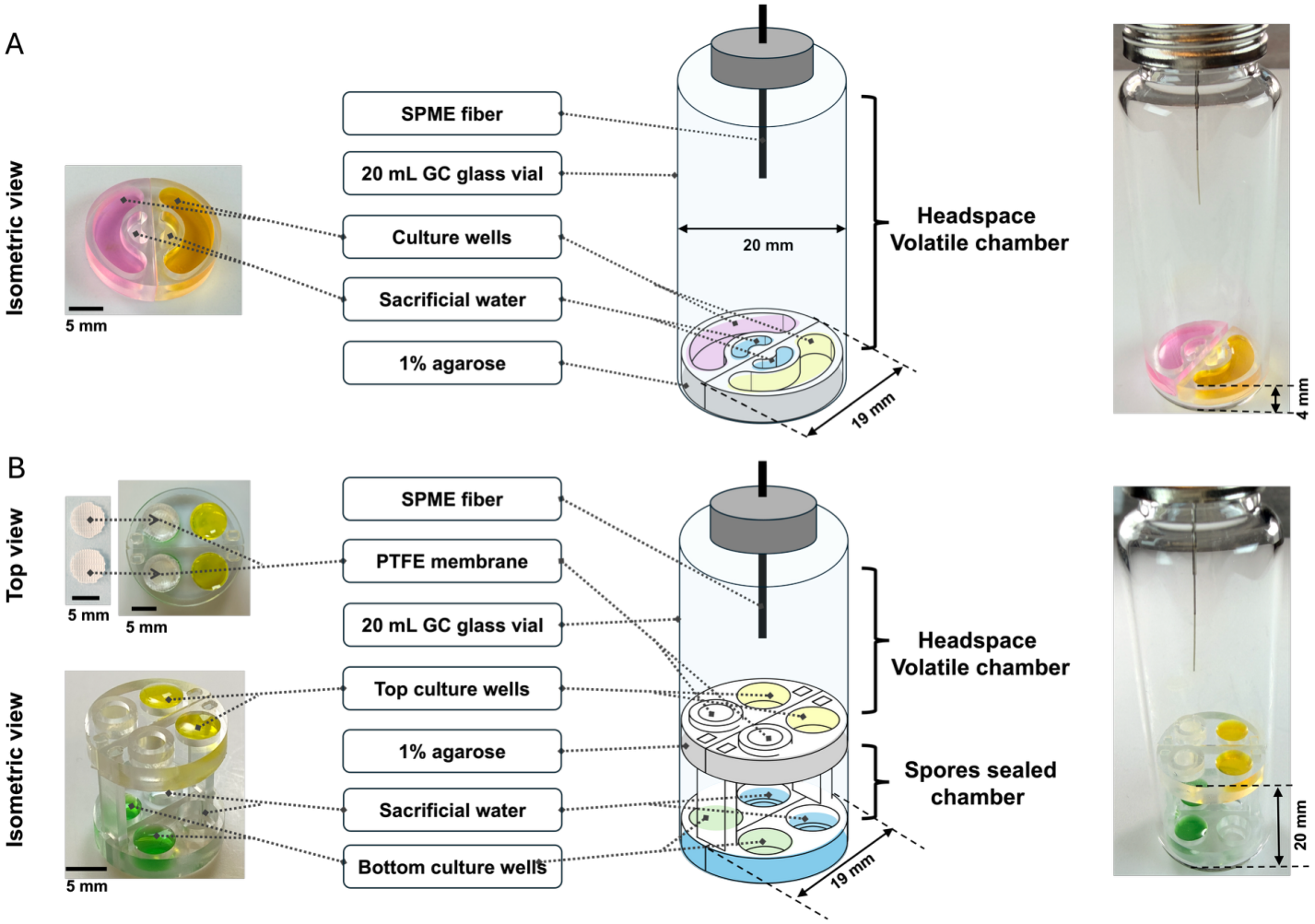
SPME GC-MS compatible mVOC co-culture device schematics.**(A)** An overview of the single-layer mVOC co-culture device showing isometric views of a polystyrene co-culture platform (left), a device schematic (middle), and a photo of the fully assembled device (right). The culture platform is designed as two halves to fit in a 20 mL GC-glass vial; 1% agarose is applied to the bottom of the vial to secure the culture platform; sacrificial water is added to the small wells to reduce evaporation. **(B)** An overview of the double-layer mVOC co-culture device, including isometric and top views of polystyrene co-culture platform (left), an isometric device schematic (middle) and a photo of the fully assembled device (right). The top and bottom culture platforms are designed as two halves to fit in a 20 mL GC-glass vial; PTFE membranes are inserted into the two stepped holes in the top culture layer with membrane pore sizes sufficiently small to prevent spore passage; 1% agarose seals the interface between the plastic and glass vial to further prevent spore migration; sacrificial water is added to the two wells in the bottom culture layer to reduce evaporation. SPME: solid phase microextraction; PTFE: polytetrafluoroethylene. Detailed device dimensions are provided in Figures S1 and S2.

### Device assembly

Prior to assembly, 20 mL glass GC-MS vials (Supelco, St. Louis, MO) were ultrasonicated with 70% ethanol for 30 min and autoclaved at 121 °C for 30 min on a gravity cycle. Metal caps with a rubber septum (Supelco, St. Louis, MO) for the vials were also sonicated in 70% ethanol for 30 min, dried with compressed air and UV-sterilized for 20 min. The single-layer co-culture platform comprises two complementary semicircular components (left and right halves), which were joined at the bottom of the 20 mL glass vials during assembly (Figure 1A, Figures S1A and S2A). Sterile tweezers were used for handling the co-culture platforms. A 100 μL solution of 1% (m/v) agarose was dispensed around the perimeter of the co-culture platform, adhering the platform to the 20 mL glass vial.

The double-layer co-culture platform consists of two culture layers each comprising two semicircular halves, two supports, two membrane holders, and two circularly trimmed PTFE membranes (Figure 1B, Figures S1B and S2B). Each semicircular half of the culture platform contains one culture well and a stepped hole with a 6.1 mm diameter in the upper section and 4 mm diameter in the lower section. The top culture layer was prepared prior to device assembly: a PTFE membrane was placed on the lower section of the through-hole in the top culture platform, secured with a membrane holder. The membrane allows diffusion of mVOCs while preventing the passage of airborne spores. The bottom layer was positioned at the bottom of the 20 mL glass vial, followed by installation of the supports (Figure S1C). Sacrificial water (180 μL) was added to the through-hole of the bottom platform to prevent evaporation of culture media. Subsequently, the membrane-secured top culture platform was installed (Figure S1C). A 180 μL solution of 1% (m/v) agarose was dispensed to the perimeter of the top platform, forming a seal between the platform and the vial wall. This step compartmentalized the upper and lower chambers, further preventing airborne spore transfer.

### P. aeruginosa and A. fumigatus microbial culture and media

The strains of *P. aeruginosa* PAO1, *P. aeruginosa* PA14, and *A. fumigatus* 293 were gifts from Dr. Nancy Keller, Medical Microbiology & Immunology and Plant Pathology, University of Wisconsin - Madison. *A. fumigatus* strain SRRC 1607 was obtained from Dr. Geromy Moore, Southern Regional Research Laboratories, U. S. Department of Agriculture, New Orleans, Louisiana, USA. *P. aeruginosa* was cultured on LB agar (Lennox) (Sigma-Aldrich, St. Louis, MO) and incubated at 37 °C overnight. Liquid cultures were prepared by inoculating an individual colony from the agar plate in 20 mL of LB broth (Lennox) (Sigma-Aldrich, St. Louis, MO) and incubating on an orbital shaker (180 rpm) at 37 °C overnight. *A. fumigatus* was cultured on Potato Dextrose Agar (PDA) (Difco, Sparks, MD) and incubated for 72 h at 37 °C. \ Fungal spores were washed from agar plates using 5 mL of 0.1% (v/v) Tween 80 (Thermo Scientific™, Rockford, IL). Prior to inoculating the device, *P. aeruginosa* liquid culture was adjusted to 0.03 optical density (OD600) in LB broth. *A. fumigatus* spore suspension was diluted to 2×10^6^ spores/mL in PDA.

### VOC addition

Prior to inoculation, 0.8 mg/mL (v/v) and 8 mg/mL (v/v) of either 1-octen-3-ol (Sigma-Aldrich, St. Louis, MO), isopentyl alcohol (Sigma-Aldrich, St. Louis, MO) or both were supplemented in PDA with 0.8% (v/v) Tween 80. The VOC-supplemented PDA (65 μL) was added to both culture wells in the bottom layer of the device, and sterile PDA with 0.8% (v/v) Tween 80 was used for vehicle control. Both top culture wells were inoculated with 65 μL of *P. aeruginosa* liquid culture (Figure 2A). The vials were capped and secured with parafilm. Vials were then placed in a pipette tip box, which was placed in a 37 °C incubator for 24 h to allow microbial growth, mVOC production, and diffusion before bacterial growth assessments.

**Figure 2.**
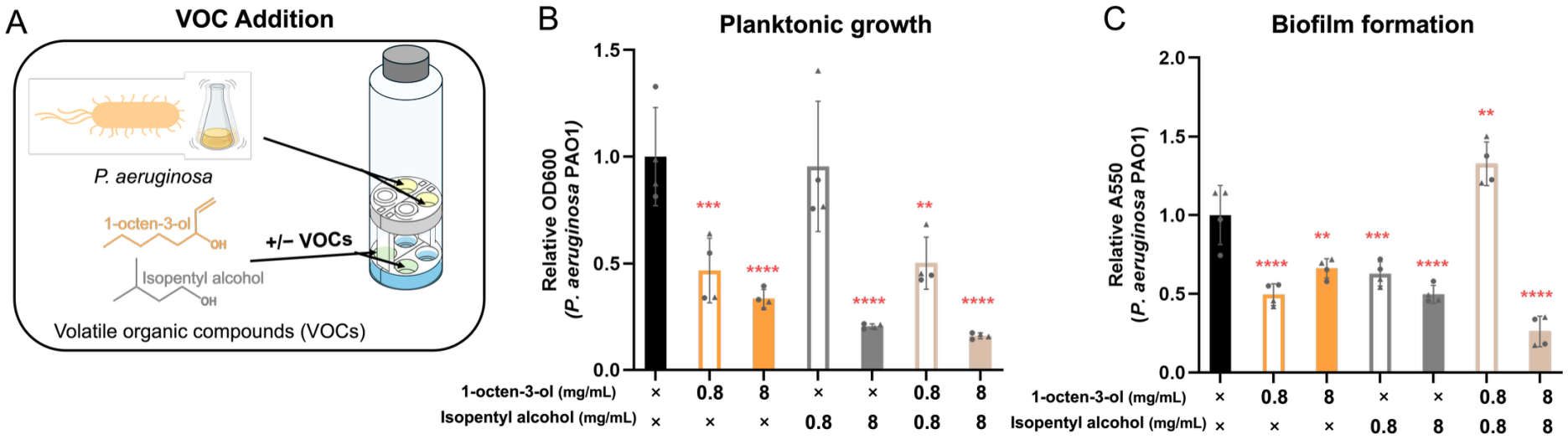
Diffusion of mVOCs across the membrane induces physiological changes of *P. aeruginosa* PAO1.**(A)** Schematic of mVOC-addition experimental conditions showing mVOC addition to bottom wells and *P. aeruginosa* cultures in top wells. **(B)** Planktonic growth of *P. aeruginosa* PAO1, after 24 h exposure to two different doses (0.8 mg/mL and 8 mg/mL) of 1-octen-3-ol and/or isopentyl alcohol. **(C)** Biofilm formation determined by crystal violet staining of *P. aeruginosa* PAO1 after 24 h exposure to two different doses (0.8 mg/mL and 8 mg/mL) of 1-octen-3-ol and/or isopentyl alcohol. Mean values were calculated for four samples originating from two culture wells in two independent devices; data points with the same shape indicate samples from the same device. The mean values were normalized to the vehicle control (black bar). The error bars represent the standard error of the mean. Statistical analysis was performed comparing each condition to the vehicle control using one-way ANOVA with Tukey’s multiple comparisons test. Statistical differences are shown with red asterisks (no asterisk, p-value > 0.05; *, p-value < 0.05; **, p-value < 0.01; ***, p-value < 0.001; ****, p-value < 0.0001).

### Co-culture inoculation

Both bottom culture wells were inoculated with either 65 μL of *A. fumigatus* spore suspension in PDA or 65 μL of sterile PDA. The top culture wells were filled with either 65 μL of *P. aeruginosa* liquid culture or 65 μL of sterile LB liquid medium. The vials were capped and parafilmed. Vials were then placed in a pipette tip box, which was placed in a 37 °C incubator for 24 h to allow microbial growth and mVOC production before sampling by HS-SPME followed by GC×GC-TOFMS data collection.

### Planktonic growth and biofilm development

After 24 h incubation, 60 μL of *P. aeruginosa* liquid culture from each culture well was aspirated and transferred to a 96-well plate. Subsequently, 40 μL of 1× phosphate-buffered saline (PBS) (pH 7.4, Gibco™, Waltham, MA) was added to each well to bring the final volume to 100 μL. Optical density (OD600) was measured to assess the planktonic growth of *P. aeruginosa*. Upon completion of OD600 measurement, the top culture wells were washed with 70 μL 1× PBS three times to remove any loosely bound bacterial cells and medium. Once the culture wells were air-dried, 70 μL of 0.1% solution of crystal violet was added to the wells and incubated with the biofilm at room temperature for 30 min. After incubation, the dye solution was removed, and the biofilms were washed three times with 1× PBS to remove free dye. A solution (70 μL) of 95% acetic acid was added to the wells, incubated for 5 min to solubilize crystal violet, and transferred to a 384-well plate. Biofilm growth was assessed and quantified by absorbance at 590 nm. A Cytation 5 Cell Imaging Multi-Mode Reader (Aligent: BioTek, Winooski, VT) was used for all the optical density and absorbance measurements. GraphPad Prism 6 software (La Jolla, CA) was used for statistical analysis. All tests were performed using one-way ANOVA with Tukey’s multiple comparisons test.

### HS-SPME-GC×GC-TOFMS Conditions

The GC×GC-TOFMS data were acquired using a Pegasus BT 4D GC×GC-TOFMS system equipped with a quad-jet thermal modulator (LECO, St. Joseph, MI, USA) and an Agilent 7890 GC (Agilent Technologies, Palo Alto, CA, USA). Sample extraction was performed using an L-PAL3 autosampler (LECO, St. Joseph, MI, USA), equipped with a temperature-controlled incubation unit and robotic arm. After the sample vials were loaded into the autosampler, the programmed incubation, headspace extraction, injection, and data acquisition sequence was performed automatically without manual handling between these steps.

A “set” consisted of one sample from each of the four sample classes (M, PA, AF, and CC). The collection sequence within each set was randomized. Six sets were collected, with two sets analyzed per collection day, resulting in six biological replicates for each sample class. Each biological replicate was prepared in a separate SPME vial and analyzed by a single GC×GC-TOFMS injection. Prior to transferring to the autosampler, all samples were incubated at 37 °C in a standard incubator used for microbial culture. Samples were then transferred to the temperature-controlled incubation of the L-PAL autosampler maintained at 37 °C while awaiting the programmed extraction and analysis sequence.

The headspace of each sample was extracted using a divinylbenzene/carboxen/poly-dimethylsiloxane (DVB/CAR/PDMS) SPME fiber with a 50/30 μm coating thickness (Supelco, Bellefonte, PA, USA). The same fiber was used throughout the study. At the beginning of each collection day, the SPME fiber was conditioned in the GC inlet at 250 °C for 1 h. The SPME workflow was standardized to maintain a consistent 1 h interval between sample injections. Prior to each extraction, the fiber was conditioned at 250 °C for 5 min. For headspace extraction, the autosampler inserted the SPME fiber assembly into the sample vial and exposed the fiber to the vial headspace at 37 °C for 30 min to extract volatile analytes. The fiber was then retracted into its protective needle and transferred by the autosampler to the GC inlet, where it was thermally desorbed under splitless conditions at 250 °C for 5 min for subsequent GC×GC-TOFMS analysis. Between successive extractions and chromatographic runs, the fiber was reconditioned at 250 °C for 20 min.

Chromatographic separations were performed using a non-polar Rxi-5Sil MS column as the first dimension (^1^D) column (60 m length × 0.25 mm i.d. × 0.25 μm film) coupled with a midpolar Rtx-200 column as the second dimension (^2^D) column (2.0 m length × 0.18 mm i.d. × 0.20 μm film). The ^1^D oven temperature program began at 40 °C and was held for 5 min, followed by a ramp of 8

°C/min to 250 °C, and held for 10 min. The ^2^D oven and thermal modulator were maintained at temperature offsets of +5 °C and +18 °C, respectively, using the same temperature program. Ultrahigh purity helium (Grade 5; Praxair, Seattle, WA, USA) was used as the carrier gas at a constant flow rate of 1.4 mL/min. The modulation period was 3 s. The transfer line and ion source temperatures were maintained at 285 °C and 225 °C, respectively. Mass spectra were acquired over an *m/z* range of 40-334 at an acquisition rate of 100 Hz using 70 eV electron ionization following a 10 s acquisition delay.

### HS-SPME-GC×GC-TOFMS Data Analysis

All HS-SPME-GC×GC-TOFMS data was processed in MATLAB R2021b (MathWorks, Natick, MA, USA). The GC×GC chromatograms were baseline corrected on a per *m/z* basis using a rolling-ball algorithm and cropped to a retention time window of 15-35 min in ^1^D and 0.5-3 s in ^2^D to exclude stationary phase overloading peaks and focus on chromatographic regions containing analytes of interest.^19,20^ Chromatograms were normalized within each sample class across six biological replicates. ^19,20^ The dataset comprised four sample classes with six biological replicates each (24 samples total), which were subjected to tile-based F-ratio analysis using an in-house MATLAB algorithm as previously described.^32–34^ A tile size of 15 s (five modulations) in ^1^D and 200 ms (20 spectra) in ^2^D was used to match typical GC×GC peak dimensions. An F-ratio threshold of 0 to ensure discovery of all detectable analyte hits, a signal-to-noise ratio (*S/N*) threshold of 10, and a minimum of 3 *m/z* per tile were applied to generate the initial hit list.

For each hit, the *m/z* corresponding to the maximum F-ratio was selected, and signals within each hit tile were summed across the ^2^D dimension to generate a ^1^D chromatographic profile and likewise across the ^1^D dimension to generate a ^2^D chromatographic profile. The resulting profiles were examined to verify the presence of a chromatographic peak and remove artifacts. Since both ^1^D and ^2^D profiles in principle provided the same analyte peak area, the ^2^D profiles of the analyte hits were subsequently subjected to secondary baseline correction to minimize residual baseline variation among replicates.^31^ The resulting baseline-corrected summed ^2^D peak area signals were used to perform one-way analysis of variance (ANOVA) across the four sample classes at the 99% confidence level (CL) (p < 0.01). For each analyte hit, the baseline-corrected ^2^D peak area signals were also averaged across the six biological replicates within each sample class, and the corresponding standard deviations were calculated. For analytes passing ANOVA, the mean signal across the six biological replicates for each sample class was normalized to the maximum mean signal across the four sample classes. PCA was performed on the resulting normalized class-mean signals in MATLAB using economy-size singular value decomposition via the built-in svd function. Chromatographic profiles in both dimensions were used for subsequent visualization.

To further characterize relationships between sample classes, six pairwise t-tests were performed at 95% CL (p < 0.05) for all pairwise class comparisons: M vs PA, M vs AF, M vs CC, PA vs AF, PA vs CC, and AF vs CC. Each analyte hit was assigned a six-digit ternary code (−1, 0, or 1) based on the outcomes of the six pairwise t-tests, with each digit corresponding to a pairwise comparison in the order listed above. A value of 1 indicates a significantly higher mean signal in the first class of each pair, −1 indicates a significantly higher mean signal in the second class, and 0 indicates no significant difference between sample classes. Analyte hits sharing the same recurring ternary-code pattern were assigned to ternary-code groups, and groups exhibiting similar class-dependent response patterns were further organized into broader Categories. Singly occurring ternary-code patterns were retained as unique patterns. The ternary-code classifications were subsequently used to color and shape the PCA score plots, enabling visualization of relationships among analyte response patterns.

## RESULTS AND DISCUSSION

### mVOC co-culture device design

To isolate and characterize mVOC-mediated communication between organisms, we engineered two microscale microbial co-culture models that consist of a unique culture platform enclosed in a standard 20 mL GC-MS glass vial, offering a broad range of microbial compatibility and allowing direct insertion of a SPME fiber for easy integration into GC-MS workflows. Importantly, as described in more detail in the Methods section, the GC-MS vials were loaded into a standard autosampler with a robotic arm that facilitated automated SPME extraction, transfer of the SPME fiber to the GC inlet, and analysis of extracted analytes. Volatile compounds collected in these devices are amenable to efficient and high-throughput sampling via HS-SPME-GC×GC-TOFMS. The two devices are simple to fabricate, implement, and interrogate, supporting different microbial systems. The single-layer mVOC co-culture device is designed to accommodate non-spore-forming microorganisms, comprising two physically separated culture wells (Figure 1A). This model enables volatile metabolite exchange while preventing physical interactions between microbial species.

To investigate mVOC crosstalk between non-spore-forming and spore-forming microbes, we designed a double-layer mVOC co-culture device (Figure 1B). This device contains two chambers connected by shared headspace via PTFE membranes, which have a pore size (1 μm) smaller than most airborne spores,^38^ thus preventing spore diffusion across chambers. Additionally, 1% agarose was used as a seal between the perimeter of the top-culture-layer and the walls of the glass vial to ensure complete physical separation of the two chambers. Materials incorporated in the system are both non-volatile and inert. Instead of gluing the PTFE membrane to the device, it was secured by sandwiching between the top culture platform and a membrane holder (Figure S1B). To evaluate the membrane and double-layer structure in preventing spore diffusion, the known spore-forming fungus *A. fumigatus* was inoculated in the bottom layer, and the top culture wells were filled with sterile PDA to capture potential spore contamination. No fungal growth of *A. fumigatus* was observed in the spore-capture medium after 2 days of incubation, as further verified by microscopy, confirming a physical separation between the two chambers (Figure S3). This unique and novel double-layer feature facilitates broad microbial compatibility with the bottom chamber accommodating diverse microbes and the upper chamber designed for non-spore-forming microbes, while preventing spore-derived physical interactions. Moreover, the design of four separate culture wells (Figures 1B and 2B) supports cultures of up to four microbial species, allowing the investigation of mVOC interaction across multiple levels, including inter- and intra-kingdom communication. In the subsequent studies, we employed the double-layer device to interrogate the interkingdom mVOC communications between two microbial pathogens, a non-spore-forming bacterium and a spore-forming fungus.

### Effects of mVOCs produced by *A. fumigatus* on *P. aeruginosa*

We selected *A. fumigatus* and *P. aeruginosa* as mVOC producing organisms to test our device. Co-infections of these pathogenic organisms are common, and volatile compounds produced by these organisms have been identified that may mediate disease-associated effects.^1–3,39^ Prior to co-cultivation of *A. fumigatus* and *P. aeruginosa*, we assessed the capability of the double-layer device in facilitating mVOC gas-phase diffusion through exogenous mVOC supplementation. Previous studies identified 1-octen-3-ol and isopentyl alcohol as the most abundant mVOCs produced by *A. fumigatus* SRRC1607 at 37°C and isopentyl alcohol as the main mVOC produced by *A. fumigatus* 293.^40,41^ Additionally, direct liquid addition of 1-octen-3-ol and volatile addition of isopentyl alcohol were shown previously to disrupt bacterial cell membrane integrity and impair growth.^42,43^ Thus, to evaluate mVOC diffusion within the device, we exogenously supplemented these compounds to the lower chamber of the device at two different concentrations (8 mg/mL and 0.8 mg/mL) and inoculated *P. aeruginosa* in the upper culture chamber (Figure 2A). We found that these volatile compounds induced physiological changes of *P. aeruginosa*, which confirms efficient headspace-mediated mVOCs transfer across the PTFE membrane within the device. The planktonic growth of *P. aeruginosa* strains PAO1 and PA14 was attenuated by 8 mg/mL of 1-octen-3-ol, isopentyl alcohol, and a mixture of the two compounds (Figures 2B and S4A), demonstrating the antibacterial activity of these compounds against *P. aeruginosa* in volatile forms. Supplementation of 0.8 mg/mL of 1-octen-3-ol and the mixture suppressed planktonic growth of *P. aeruginosa* PAO1 but not *P. aeruginosa* PA14, while 0.8 mg/mL of isopentyl alcohol did not impact growth of either *P. aeruginosa* (Figure 2B and S3A). These results imply that *P. aeruginosa* PAO1 is more susceptible to 1-octen-3-ol than *P. aeruginosa* PA14, and both strains are resistant to low dosage of isopentyl alcohol volatiles. Notably, 0.8 mg/mL of 1-octen-3-ol is 2.5-fold lower than the minimal inhibitory concentrations (MIC) of liquid-phase addition reported by Xiong et al.^42^ The observed effect at lower dosages in our experiment may be due to the sealed environment of the device enabling accumulation of 1-octen-3-ol in the vapor-phase, thus sustaining bacterial exposure and/or increasing the effective concentration relative to liquid-phase addition.

In addition to growth assessments, our double-layer device is compatible with crystal violet assays to capture biofilm formation. Biofilms are communities of microorganisms that adhere to each other and to biotic and abiotic surfaces, protecting against antibiotics and host immune attacks by a self-produced extracellular matrix of polymeric substances.^3,44^ *P. aeruginosa* is a well-known biofilm producer and previous studies have shown that direct contact with 1-octen-3-ol resulted in reduced biofilm formation.^45^ Volatile exposure to isopentyl alcohol also resulted in reduced biofilm formation in both gram-positive and gram-negative bacteria, including *P. aeruginosa*.^46^ In agreement with these studies, we detected a reduction in biofilms for both *P. aeruginosa* strains after addition of 1-octen-3-ol, isopentyl alcohol, and their mixtures at 8 mg/mL in volatile forms (Figures 2C and S4B). Additionally, we found that the biofilm development of *P. aeruginosa* PAO1 was reduced upon exposure to 0.8 mg/mL isopentyl alcohol while that of *P. aeruginosa* PA14 remained unaffected (Figure 2C, S4B), indicating strain level variation against anti-biofilm mVOCs. Interestingly, we found that the mixture of 0.8 mg/mL 1-octen-3-ol and isopentyl alcohol promoted *P. aeruginosa* PAO1 biofilm formation (Figure 2C). This observed stimulatory effect resembles the biofilm development detected in *P. aeruginosa* PAO1 under sub-MIC antibiotic treatment, which was proposed as a survival strategy against antibiotic stress.^47^ Together, these biological assays demonstrate the versatility of our double-layer device across a wide range of mVOC-mediated studies.

After validating our device using exogenously supplemented volatile compounds, we evaluated its ability to capture the exchange of naturally produced mVOCs between microbial co-cultures. We tested whether co-cultivation of *A. fumigatus* and *P. aeruginosa* strains would induce similar physiological changes to *P. aeruginosa* to those observed following exposure to purified volatile compounds produced by *A. fumigatus*. Unexpectedly, there was no detectable effect on the planktonic growth and biofilm formation of *P. aeruginosa* strains in co-cultures with *A. fumigatus* compared to a media control (Figure S5). One plausible explanation is that *A. fumigatus* produced low levels of 1-octen-3-ol and isopentyl alcohol under the tested conditions, which were insufficient to induce biological responses measured in this study. Another possibility is that the presence of other volatiles or their combined effects altered the overall impact of these inhibitory molecules. Alternatively, the mutual mVOC interactions in the co-culture could potentially be dominated by *P. aeruginosa*, thereby masking the effects of A. *fumigatus* mVOCs. The effects of *P. aeruginosa*-derived mVOCs on *A. fumigatus* are presently unclear with previous studies reporting opposing findings.^48–50^ Thus, further studies are needed to understand the underlying biological mechanisms responsible for mVOC-mediated communications between these opportunistic pathogens.

### HS-SPME-GC×GC-TOFMS Analysis

To characterize the volatile metabolome associated with microbial co-culture, headspace samples from four sample classes, including media blank (M), *P. aeruginosa* PAO1 monoculture (PA), *A. fumigatus* SRRC1607 monoculture (AF), and co-culture of PA and AF (CC), were analyzed by HS-SPME-GC×GC-TOFMS. The total ion current (TIC) chromatograms demonstrate the strong separation capability of GC×GC-TOFMS (Figure 3), resolving hundreds of volatile compounds across the 2D chromatographic space. Despite the complexity of the data, sample class-dependent differences are visually apparent when comparing the raw TIC chromatograms. For example, the region highlighted by the red box contains an analyte detected in the M and AF chromatograms but absent from the PA and CC, suggesting that this compound is depleted by *P. aeruginosa* in both monoculture and co-culture conditions (Figure 3). However, this comparison also illustrates the analytical challenge of interpreting GC×GC-TOFMS data, where each chromatogram contains hundreds of features with subtle and overlapping class differences.

**Figure 3.**
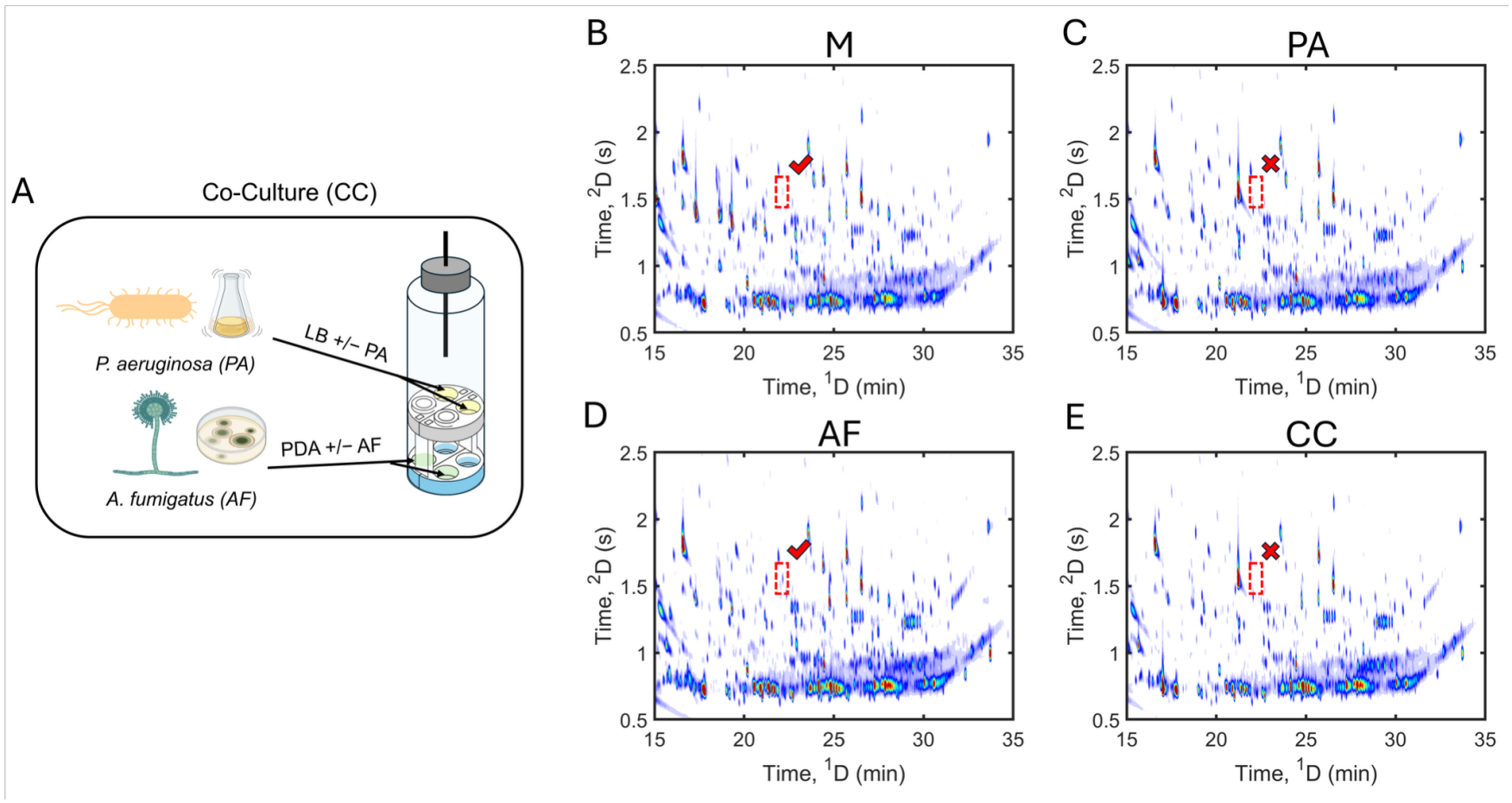
GC×GC-TOFMS identifies volatile metabolites present in co-culture device experiments. **(A)** Schematic of co-culture experimental conditions for GC×GC-TOFMS showing *A. fumigatus* SRRC 1607 cultures in bottom wells and *P. aeruginosa* PAO1 cultures in top wells. **(B–E)** Total ion current (TIC) chromatograms of one representative replicate for each sample class: **(B)** M (Media), **(C)** PA (*P. aeruginosa* PAO1), **(D)** AF (*A. fumigatus* SRRC 1607), and **(E)** CC (co-culture of PA and AF). These chromatograms are representative of six replicates per sample class. The red box indicates an example analyte detected in PA and CC but not detected in AF and Media.

To address this challenge, tile-based F-ratio analysis was performed to identify analytes that differed across at least two of the four sample classes, resulting in a total of 198 analyte hits after artifact removal.^19,20^ These hits were further evaluated by one-way ANOVA across the four sample classes at the 99% CL, which yielded 104 hits passing the significance threshold with p-values <0.01 (Figure S6). Thus, for each of the 104 hits, the mean signal differed significantly between at least two of the four sample classes. The chromatographic profiles and signal intensity for statistically significant hits were first evaluated qualitatively across sample classes (Figure S7-S12). Four representative hits illustrating distinct signal patterns are shown in detail (Figure 4). For example, analysis of Hit 2 shows that it is present in PA and CC, but absent from media and AF, consistent with a metabolite produced by PA that is not affected by AF in co-culture. Hit 41 is representative of metabolites that have elevated levels in AF, but the levels are lower in CC relative to AF monoculture, suggesting that PA prevents accumulation of this AF-associated metabolite in co-culture and indicating a potential interspecies interaction. This qualitative analysis also reveals volatile compounds that are consumed by AF and PA. For example, Hit 4 is present in the media but depleted in both PA and CC, consistent with a media-derived compound that is utilized by *P. aeruginosa*. Hit 10 is also present in media but is depleted across the sample classes to different extents, with the lowest levels in CC, moderate levels in PA, and relatively higher levels in AF. This pattern suggests utilization by both organisms, with more extensive consumption in the co-culture condition. Together, these four representative analytes demonstrate that the 104 statistically significant hits are not a monolithic group, motivating a systematic approach to classify all 104 hits by pattern rather than examining them one at a time.

**Figure 4.**
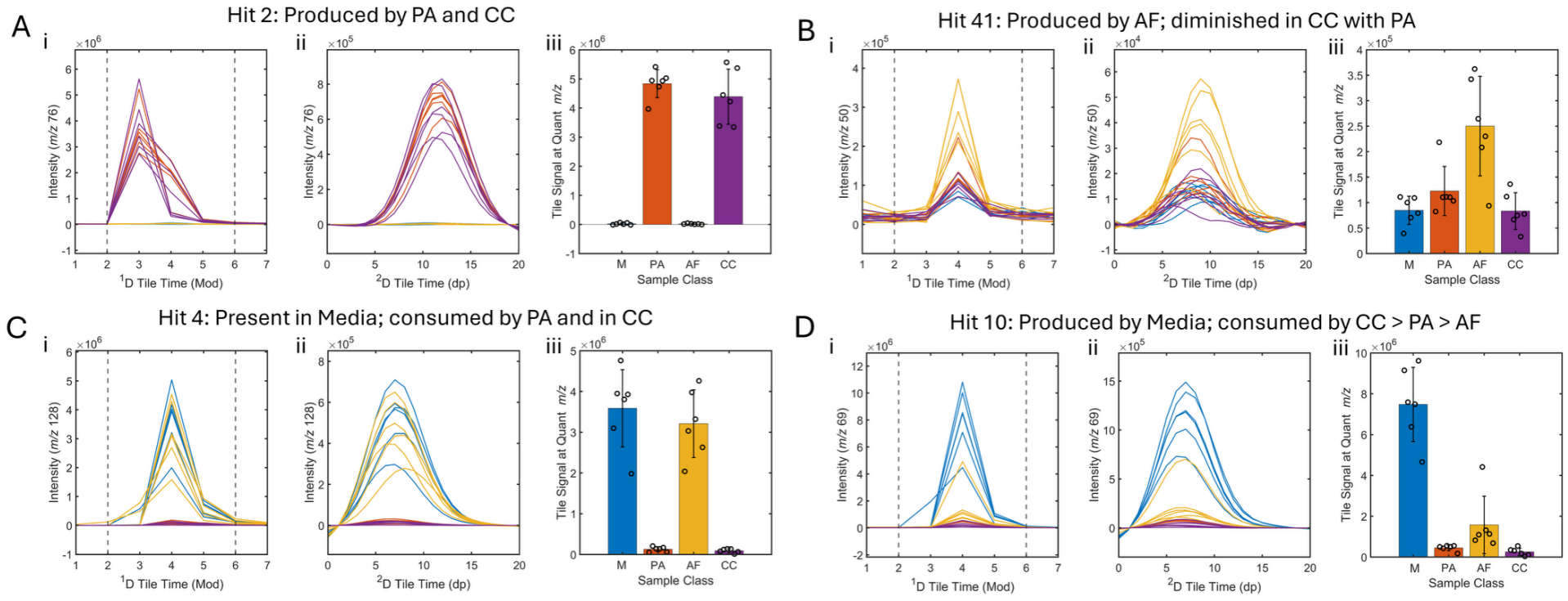
Distinctive peak profiles and signal patterns for volatile metabolites revealed from GC×GC-TOFMS comparisons across sample classes. Representative hit profiles and signal patterns of **(A)** an analyte produced by PA and CC (Hit 2), **(B)** an analyte produced by AF (Hit 41), **(C)** an analyte present in media and consumed by CC and PA (Hit 4), and **(D)** an analyte present in media and consumed in CC, PA, and AF (Hit 10). Colors of chromatograms correspond to sample class: blue (M), red (PA), orange (AF), and purple (CC). Each hit shows (i) ^1^D chromatographic profile (signal summed across ^2^D), (ii) ^2^D chromatographic profile (signal summed across ^1^D), and (iii) signal profile patterns at selective *m/z*. Data points in A-D (iii) represent each of six replicate signals; error bars in A-D (iii) represent the standard error of the mean.

As an initial, unsupervised approach to visualizing groupings among the 104 ANOVA-significant analytes, PCA was performed on the mean, max-normalized signal profiles (Figure 5). This analysis shows clear separation of the four representative hits from Figure 4 based on their principal components, consistent with their distinct qualitative patterns across sample classes. However, the 104 significant analytes did not separate into clearly defined clusters, initially limiting the insights from this analysis. In addition, PCA alone does not indicate which class-pair comparisons drive an analyte’s position or assign it to a discrete, biologically meaningful group. These limitations motivated the use of a ternary-code classification scheme, derived from pairwise t-tests, as a complementary, supervised approach to classify the 104 hits into discrete, interpretable pattern groups.

**Figure 5.**
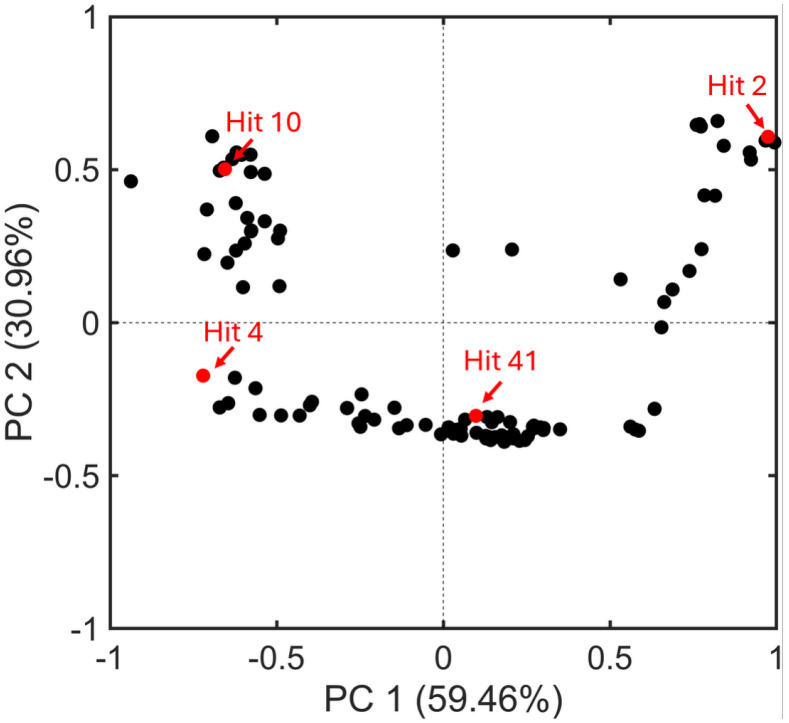
PCA scores plot of signal profile patterns per Fig. 4A-D(iii) shows the continuous distribution of 104 significant analytes from ANOVA analysis (99% CL). Each signal was normalized to the maximum signal across the four sample classes, resulting in values ranging from 0 to 1. Four representative analytes with signal profiles shown in Figure 4 are highlighted in red.

For these 104 hits, six pairwise t-tests (95% CL) were performed across all class-pair combinations (M-PA, M-AF, M-CC, PA-AF, PA-CC, AF-CC), and each analyte was assigned a six-digit ternary code (1, −1, or 0 per comparison) to indicate whether its signal was significantly higher, significantly lower, or not significantly different between each pair of classes (Tables S1 and S2). This ternary code is the key output that links quantitative statistical testing to biologically interpretable “production” or “consumption” patterns for each hit and underlies the proposed classification. With six ternary-code positions, each taking a value of −1, 0, or 1, there are 729 patterns theoretically possible. Under an equal-probability model, the probability that a specific ternary-code pattern occurs exactly twice among 104 analytes is approximately 0.0088. To assess repeated patterns across all possible codes, a Monte Carlo simulation was performed (and compared to probability theory) in which 104 hits were independently assigned with equal probability to the 729 possible six-digit ternary codes (see detailed description in Supporting Information). This process was repeated 10^6^ times. Among the 96,982,390 distinct code patterns observed across 10^6^ simulations, 93.09% occurred once, 6.59% twice, 0.307% three times, 0.0106% four times, 0.000278% five times, and only six patterns occurred six times, with none occurring seven or more times (Figure S13). For subsequent classification, patterns occurring in two or more analytes were treated as recurring patterns. By applying this criterion, we identified 15 distinct ternary-code groups among the analytes with a recurring pattern, labeled A through O (Figure 6 and Table 1). Out of the 104 analytes, 88 hits are categorized into these 15 groups (Figures S7-S11), which represent a range of distinct metabolic relationships among the sample classes. These groups capture the qualitative patterns previously noted, including compounds produced exclusively by one organism and preserved in co-culture, compounds produced by one organism but suppressed in co-culture, and media-derived compounds consumed to varying degrees by PA, AF, or both. The remaining 16 hits have ternary-code patterns that only occur once (Figure S12).

**Table 1.** Compound Categories, corresponding ternary-code groups, number of hits in each group, six-digit ternary codes, and metabolite descriptions for the 88 analytes exhibiting recurring ternary-code patterns and four additional metabolites with unique ternary codes assigned to Category 6.

| **Category** | **Group** | **# Hits** | **M vs. PA** | **M vs. AF** | **M vs. CC** | **PA vs. AF** | **PA vs. CC** | **AF vs. CC** | **Metabolite Description** |
|---|---|---|---|---|---|---|---|---|---|
| **1** | A | 12 | -1 | 0 | -1 | 1 | 0 | -1 | Produced in PA and CC |
|  | H | 4 | -1 | 0 | -1 | 1 | 1 | -1 |  |
| **2** | B | 12 | 0 | -1 | 0 | -1 | 0 | 1 | Produced in AF;<br>Production suppressed or consumed in CC |
|  | F | 6 | -1 | -1 | 0 | 0 | 1 | 1 |  |
|  | I | 4 | 0 | 0 | 0 | -1 | 0 | 1 |  |
|  | K | 3 | 0 | -1 | 0 | 0 | 0 | 1 |  |
|  | M | 2 | 0 | -1 | 1 | -1 | 0 | 1 |  |
| **3** | C | 12 | 1 | 0 | 1 | -1 | 0 | 1 | Present in Media;<br>Consumed in PA and CC |
|  | J | 4 | 1 | -1 | 1 | -1 | 0 | 1 |  |
|  | L | 3 | 1 | 0 | 1 | 0 | 0 | 1 |  |
| **4** | D | 9 | 1 | 1 | 1 | 0 | 0 | 0 | Present in Media;<br>Consumed in PA, AF, and CC |
|  | E | 8 | 1 | 1 | 1 | -1 | 0 | 1 |  |
|  | G | 5 | 1 | 1 | 1 | 0 | 1 | 0 |  |
|  | O | 2 | 1 | 1 | 1 | 0 | 0 | 1 |  |
| **5** | N | 2 | 0 | 1 | 1 | 1 | 1 | 0 | Present in Media;<br>Consumed in AF and CC |
| **6** | N/A | 4 | -1 | -1 | VAR* | 0 | VAR* | VAR* | Produced in PA, AF, and CC |

\*VAR indicates positions where the ternary-code digits vary among the four metabolites in Category 6.

**Figure 6.**
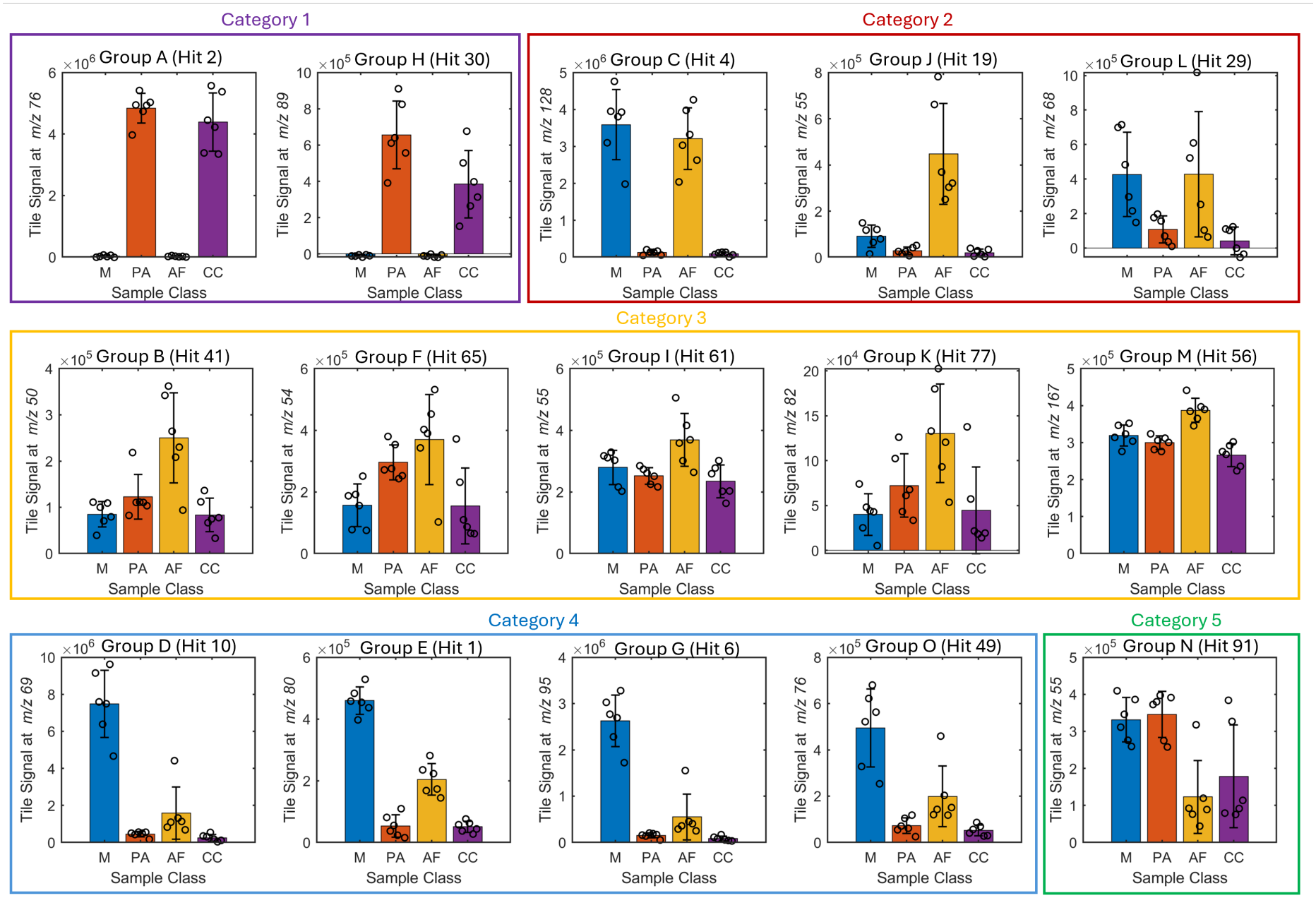
Ternary-code classification reveals 15 distinct volatile metabolite groups (A-O) with characteristic signal patterns. Signal patterns of one representative hit at selective *m/z* per ternary group are shown. Bar colors correspond to sample class: blue (M), red (PA), orange (AF), and purple (CC). Data points represent each of six replicate signals; error bars represent the standard error of the mean. The 15 groups are further organized into five Categories (Table 1) as determined by ternary-code patterns and PCA analysis.

Finally, to connect this discrete ternary classification approach to the unsupervised PCA, the assigned ternary-code groups for each analyte were mapped onto the original PCA scores plot (Figure S14). Several distinct ternary-code groups are intermixed or positioned near one another in the PCA score space, and inspection of their ternary codes showed that these groups often differ by only one or a few positions in their ternary codes. These similarities motivated further consolidation of the 15 ternary-code groups into 5 major Categories based on their similarities in ternary-group patterns and relative positions in the PCA score space (Figure 7), visualized with ellipses indicating approximate Category boundaries. The compound Categories, corresponding ternary-code groups, number of hits in each group, six-digit ternary codes, and metabolite descriptions are summarized in Table 1. Notably, once the analytes are colored by Category, the PCA axes themselves take on clear biological meaning: The PC1 axis correlates with decreasing metabolite abundance in Media from left to right, while the PC2 axis correlates with increasing metabolite abundance in AF from bottom to top.

**Figure 7.**
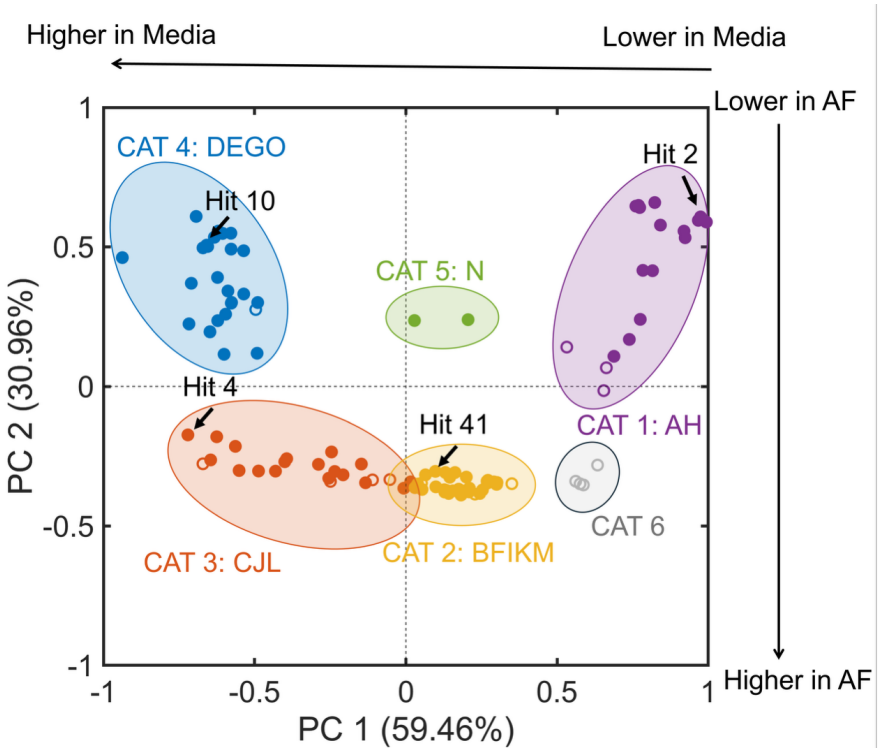
Ternary-code classification reveals clustering of 104 significant analytes into five major compound Categories (CAT), with four additional analytes grouped into CAT 6. The PCA scores plot (same as Figure 5) was constructed from signal profiles of 104 analytes passing ANOVA at the 99% CL. Each signal was normalized to the maximum signal across the four sample classes, resulting in values ranging from 0 to 1. Colors represent the compound Categories assigned based on the ternary-code groups, as summarized in Table 1. Ellipses indicate visual Category boundaries. The arrow along the PC1 axis indicates analyte abundance in media decreasing from left to right; the arrow along PC2 axis indicates analyte abundance in AF increasing from bottom to top. Open circles represent 16 analytes with ternary codes occurring only once, color-coded by their proximity to the five defined Categories in the scores plot, except for four analytes grouped into Category 6, which are colored gray.

Of the 104 analytes, 16 had unique ternary codes occurring only once and are represented in the PCA plot by open circles. Among these analytes, 12 are positioned in close proximity to the five defined Categories and have been color-coded accordingly. Indeed, these analytes have ternary codes that differ from those of the nearby recurring groups by only one or a few positions, reflecting similar overall signal patterns across the five Categories. These differences may result from individual pairwise comparisons that did not reach statistical significance, while the overall patterns remain similar to those of the corresponding Categories. The remaining four analytes fall outside all five Category boundaries and are shown as gray open circles, representing analytes with unique, Category-independent patterns not well captured by the five dominant Categories. However, their PCA positions and signal patterns remain consistent with the biological trends represented by PC1 and PC2, suggesting that their unique ternary codes nevertheless capture biological patterns consistent with their positions in the PCA score space.

Overall, this alignment between the unsupervised PCA structure and the supervised ternary classification is an important internal validation of the classification scheme, confirming that the five defined Categories correspond to underlying, PC-resolvable gradients in media metabolite consumption and AF-associated metabolite production, and substantially improving interpretability over the initial PCA shown in Figure 5.

### mVOC profiles of mono- and co-cultures of *A. fumigatus* and *P. aeruginosa*

Next, we made tentative chemical assignments for the analytes in each Category to profile mVOCs in PA and AF monocultures and co-culture. Tentative chemical identifications for all 104 analytes, along with their ternary-code group, Category assignment, NIST library match value (MV), chemical formula, and retention times in both dimensions, are provided in Table 2. Tentative identifications were assigned using the highest NIST library match with MV ≥ 800;^31,51^ analytes with MV < 800 were reported as unknown. Our data reveal known mVOCs consistent with previous reports and also expand the known volatilomes of these microbes to include additional compounds and to provide insights into their potential roles in microbial interactions.

**Table 2.**
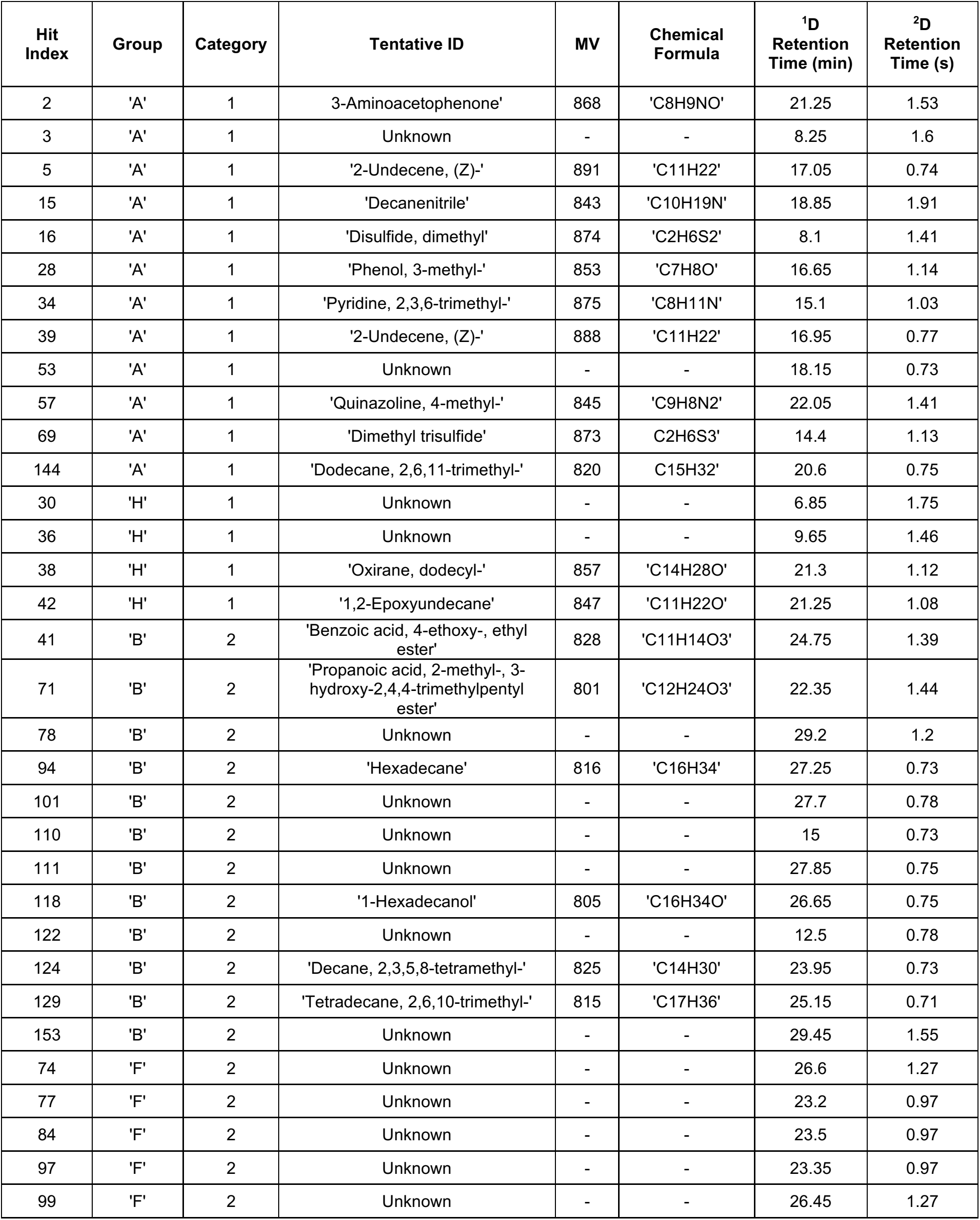
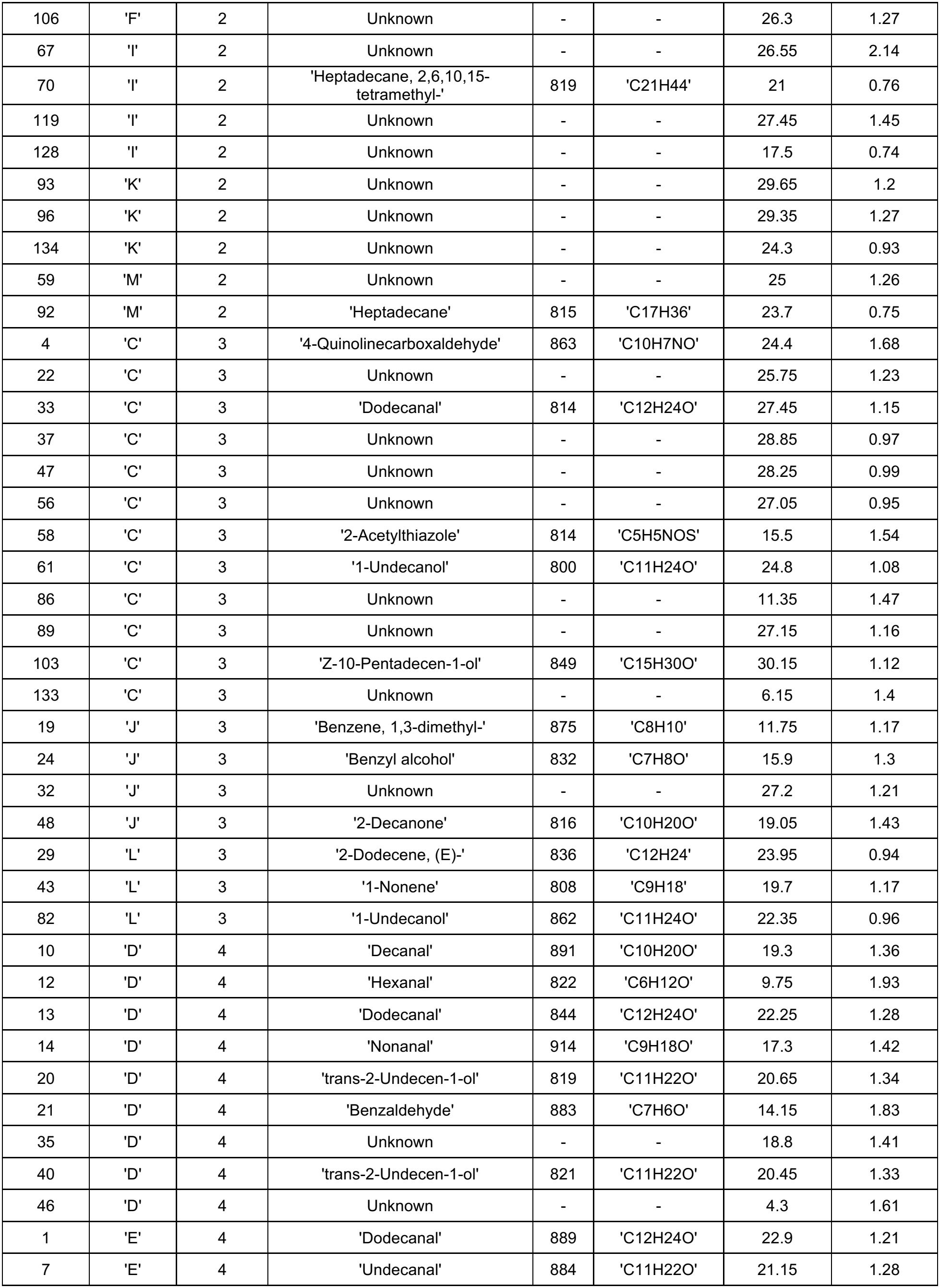
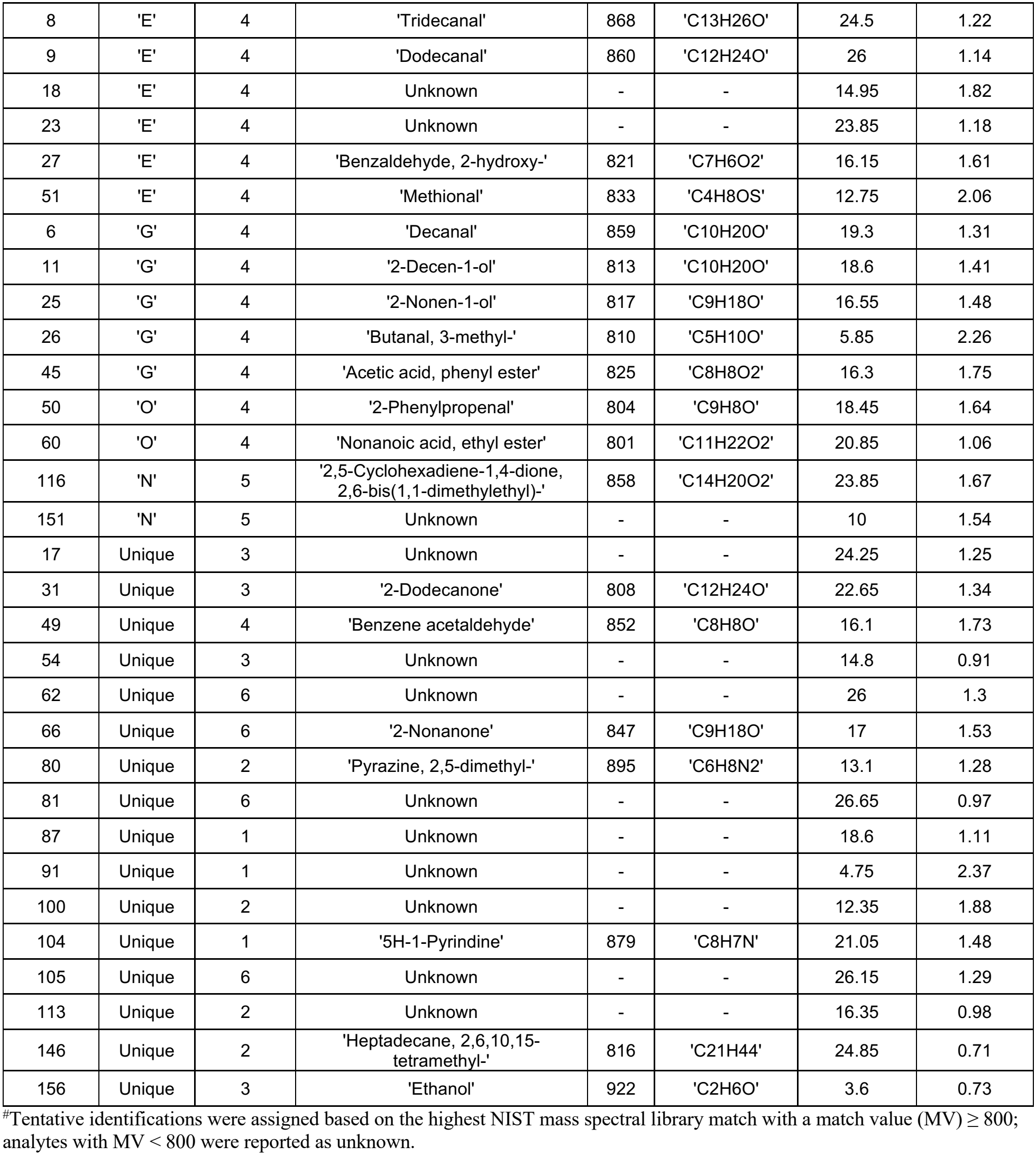
Identification and classification of the 104 analytes passing ANOVA at 99% CL. For each hit, the table lists the hit index, its ternary-code group and corresponding Category, the tentative identification from the NIST library with associated match value (MV)^#^, the chemical formula, and the retention time in both chromatographic dimensions.

| Hit Index | Group | Category | Tentative ID | MV | Chemical Formula | <sup>1</sup>D Retention Time (min) | <sup>2</sup>D Retention Time (s) |
|---|---|---|---|---|---|---|---|
| 2 | 'A' | 1 | 3-Aminoacetophenone' | 868 | 'C8H9NO' | 21.25 | 1.53 |
| 3 | 'A' | 1 | Unknown | - | - | 8.25 | 1.6 |
| 5 | 'A' | 1 | '2-Undecene, (Z)-' | 891 | 'C11H22' | 17.05 | 0.74 |
| 15 | 'A' | 1 | 'Decanenitrile' | 843 | 'C10H19N' | 18.85 | 1.91 |
| 16 | 'A' | 1 | 'Disulfide, dimethyl' | 874 | 'C2H6S2' | 8.1 | 1.41 |
| 28 | 'A' | 1 | 'Phenol, 3-methyl-' | 853 | 'C7H8O' | 16.65 | 1.14 |
| 34 | 'A' | 1 | 'Pyridine, 2,3,6-trimethyl-' | 875 | 'C8H11N' | 15.1 | 1.03 |
| 39 | 'A' | 1 | '2-Undecene, (Z)-' | 888 | 'C11H22' | 16.95 | 0.77 |
| 53 | 'A' | 1 | Unknown | - | - | 18.15 | 0.73 |
| 57 | 'A' | 1 | 'Quinazoline, 4-methyl-' | 845 | 'C9H8N2' | 22.05 | 1.41 |
| 69 | 'A' | 1 | 'Dimethyl trisulfide' | 873 | C2H6S3' | 14.4 | 1.13 |
| 144 | 'A' | 1 | 'Dodecane, 2,6,11-trimethyl-' | 820 | C15H32' | 20.6 | 0.75 |
| 30 | 'H' | 1 | Unknown | - | - | 6.85 | 1.75 |
| 36 | 'H' | 1 | Unknown | - | - | 9.65 | 1.46 |
| 38 | 'H' | 1 | 'Oxirane, dodecyl-' | 857 | 'C14H28O' | 21.3 | 1.12 |
| 42 | 'H' | 1 | '1,2-Epoxyundecane' | 847 | 'C11H22O' | 21.25 | 1.08 |
| 41 | 'B' | 2 | 'Benzoic acid, 4-ethoxy-, ethyl ester' | 828 | 'C11H14O3' | 24.75 | 1.39 |
| 71 | 'B' | 2 | 'Propanoic acid, 2-methyl-, 3-hydroxy-2,4,4-trimethylpentyl ester' | 801 | 'C12H24O3' | 22.35 | 1.44 |
| 78 | 'B' | 2 | Unknown | - | - | 29.2 | 1.2 |
| 94 | 'B' | 2 | 'Hexadecane' | 816 | 'C16H34' | 27.25 | 0.73 |
| 101 | 'B' | 2 | Unknown | - | - | 27.7 | 0.78 |
| 110 | 'B' | 2 | Unknown | - | - | 15 | 0.73 |
| 111 | 'B' | 2 | Unknown | - | - | 27.85 | 0.75 |
| 118 | 'B' | 2 | '1-Hexadecanol' | 805 | 'C16H34O' | 26.65 | 0.75 |
| 122 | 'B' | 2 | Unknown | - | - | 12.5 | 0.78 |
| 124 | 'B' | 2 | 'Decane, 2,3,5,8-tetramethyl-' | 825 | 'C14H30' | 23.95 | 0.73 |
| 129 | 'B' | 2 | 'Tetradecane, 2,6,10-trimethyl-' | 815 | 'C17H36' | 25.15 | 0.71 |
| 153 | 'B' | 2 | Unknown | - | - | 29.45 | 1.55 |
| 74 | 'F' | 2 | Unknown | - | - | 26.6 | 1.27 |
| 77 | 'F' | 2 | Unknown | - | - | 23.2 | 0.97 |
| 84 | 'F' | 2 | Unknown | - | - | 23.5 | 0.97 |
| 97 | 'F' | 2 | Unknown | - | - | 23.35 | 0.97 |
| 99 | 'F' | 2 | Unknown | - | - | 26.45 | 1.27 |
| 106 | 'F' | 2 | Unknown | - | - | 26.3 | 1.27 |
| 67 | 'I' | 2 | Unknown | - | - | 26.55 | 2.14 |
| 70 | 'I' | 2 | 'Heptadecane, 2,6,10,15-tetramethyl-' | 819 | 'C21H44' | 21 | 0.76 |
| 119 | 'I' | 2 | Unknown | - | - | 27.45 | 1.45 |
| 128 | 'I' | 2 | Unknown | - | - | 17.5 | 0.74 |
| 93 | 'K' | 2 | Unknown | - | - | 29.65 | 1.2 |
| 96 | 'K' | 2 | Unknown | - | - | 29.35 | 1.27 |
| 134 | 'K' | 2 | Unknown | - | - | 24.3 | 0.93 |
| 59 | 'M' | 2 | Unknown | - | - | 25 | 1.26 |
| 92 | 'M' | 2 | 'Heptadecane' | 815 | 'C17H36' | 23.7 | 0.75 |
| 4 | 'C' | 3 | '4-Quinolinecarboxaldehyde' | 863 | 'C10H7NO' | 24.4 | 1.68 |
| 22 | 'C' | 3 | Unknown | - | - | 25.75 | 1.23 |
| 33 | 'C' | 3 | 'Dodecanal' | 814 | 'C12H24O' | 27.45 | 1.15 |
| 37 | 'C' | 3 | Unknown | - | - | 28.85 | 0.97 |
| 47 | 'C' | 3 | Unknown | - | - | 28.25 | 0.99 |
| 56 | 'C' | 3 | Unknown | - | - | 27.05 | 0.95 |
| 58 | 'C' | 3 | '2-Acetylthiazole' | 814 | 'C5H5NOS' | 15.5 | 1.54 |
| 61 | 'C' | 3 | '1-Undecanol' | 800 | 'C11H24O' | 24.8 | 1.08 |
| 86 | 'C' | 3 | Unknown | - | - | 11.35 | 1.47 |
| 89 | 'C' | 3 | Unknown | - | - | 27.15 | 1.16 |
| 103 | 'C' | 3 | 'Z-10-Pentadecen-1-ol' | 849 | 'C15H30O' | 30.15 | 1.12 |
| 133 | 'C' | 3 | Unknown | - | - | 6.15 | 1.4 |
| 19 | 'J' | 3 | 'Benzene, 1,3-dimethyl-' | 875 | 'C8H10' | 11.75 | 1.17 |
| 24 | 'J' | 3 | 'Benzyl alcohol' | 832 | 'C7H8O' | 15.9 | 1.3 |
| 32 | 'J' | 3 | Unknown | - | - | 27.2 | 1.21 |
| 48 | 'J' | 3 | '2-Decanone' | 816 | 'C10H20O' | 19.05 | 1.43 |
| 29 | 'L' | 3 | '2-Dodecene, (E)-' | 836 | 'C12H24' | 23.95 | 0.94 |
| 43 | 'L' | 3 | '1-Nonene' | 808 | 'C9H18' | 19.7 | 1.17 |
| 82 | 'L' | 3 | '1-Undecanol' | 862 | 'C11H24O' | 22.35 | 0.96 |
| 10 | 'D' | 4 | 'Decanal' | 891 | 'C10H20O' | 19.3 | 1.36 |
| 12 | 'D' | 4 | 'Hexanal' | 822 | 'C6H12O' | 9.75 | 1.93 |
| 13 | 'D' | 4 | 'Dodecanal' | 844 | 'C12H24O' | 22.25 | 1.28 |
| 14 | 'D' | 4 | 'Nonanal' | 914 | 'C9H18O' | 17.3 | 1.42 |
| 20 | 'D' | 4 | 'trans-2-Undecen-1-ol' | 819 | 'C11H22O' | 20.65 | 1.34 |
| 21 | 'D' | 4 | 'Benzaldehyde' | 883 | 'C7H6O' | 14.15 | 1.83 |
| 35 | 'D' | 4 | Unknown | - | - | 18.8 | 1.41 |
| 40 | 'D' | 4 | 'trans-2-Undecen-1-ol' | 821 | 'C11H22O' | 20.45 | 1.33 |
| 46 | 'D' | 4 | Unknown | - | - | 4.3 | 1.61 |
| 1 | 'E' | 4 | 'Dodecanal' | 889 | 'C12H24O' | 22.9 | 1.21 |
| 7 | 'E' | 4 | 'Undecanal' | 884 | 'C11H22O' | 21.15 | 1.28 |
| 8 | 'E' | 4 | 'Tridecanal' | 868 | 'C13H26O' | 24.5 | 1.22 |
| 9 | 'E' | 4 | 'Dodecanal' | 860 | 'C12H24O' | 26 | 1.14 |
| 18 | 'E' | 4 | Unknown | - | - | 14.95 | 1.82 |
| 23 | 'E' | 4 | Unknown | - | - | 23.85 | 1.18 |
| 27 | 'E' | 4 | 'Benzaldehyde, 2-hydroxy-' | 821 | 'C7H6O2' | 16.15 | 1.61 |
| 51 | 'E' | 4 | 'Methional' | 833 | 'C4H8OS' | 12.75 | 2.06 |
| 6 | 'G' | 4 | 'Decanal' | 859 | 'C10H20O' | 19.3 | 1.31 |
| 11 | 'G' | 4 | '2-Decen-1-ol' | 813 | 'C10H20O' | 18.6 | 1.41 |
| 25 | 'G' | 4 | '2-Nonen-1-ol' | 817 | 'C9H18O' | 16.55 | 1.48 |
| 26 | 'G' | 4 | 'Butanal, 3-methyl-' | 810 | 'C5H10O' | 5.85 | 2.26 |
| 45 | 'G' | 4 | 'Acetic acid, phenyl ester' | 825 | 'C8H8O2' | 16.3 | 1.75 |
| 50 | 'O' | 4 | '2-Phenylpropenal' | 804 | 'C9H8O' | 18.45 | 1.64 |
| 60 | 'O' | 4 | 'Nonanoic acid, ethyl ester' | 801 | 'C11H22O2' | 20.85 | 1.06 |
| 116 | 'N' | 5 | '2,5-Cyclohexadiene-1,4-dione, 2,6-bis(1,1-dimethylethyl)-' | 858 | 'C14H20O2' | 23.85 | 1.67 |
| 151 | 'N' | 5 | Unknown | - | - | 10 | 1.54 |
| 17 | Unique | 3 | Unknown | - | - | 24.25 | 1.25 |
| 31 | Unique | 3 | '2-Dodecanone' | 808 | 'C12H24O' | 22.65 | 1.34 |
| 49 | Unique | 4 | 'Benzene acetaldehyde' | 852 | 'C8H8O' | 16.1 | 1.73 |
| 54 | Unique | 3 | Unknown | - | - | 14.8 | 0.91 |
| 62 | Unique | 6 | Unknown | - | - | 26 | 1.3 |
| 66 | Unique | 6 | '2-Nonanone' | 847 | 'C9H18O' | 17 | 1.53 |
| 80 | Unique | 2 | 'Pyrazine, 2,5-dimethyl-' | 895 | 'C6H8N2' | 13.1 | 1.28 |
| 81 | Unique | 6 | Unknown | - | - | 26.65 | 0.97 |
| 87 | Unique | 1 | Unknown | - | - | 18.6 | 1.11 |
| 91 | Unique | 1 | Unknown | - | - | 4.75 | 2.37 |
| 100 | Unique | 2 | Unknown | - | - | 12.35 | 1.88 |
| 104 | Unique | 1 | '5H-1-Pyrindine' | 879 | 'C8H7N' | 21.05 | 1.48 |
| 105 | Unique | 6 | Unknown | - | - | 26.15 | 1.29 |
| 113 | Unique | 2 | Unknown | - | - | 16.35 | 0.98 |
| 146 | Unique | 2 | 'Heptadecane, 2,6,10,15-tetramethyl-' | 816 | 'C21H44' | 24.85 | 0.71 |
| 156 | Unique | 3 | 'Ethanol' | 922 | 'C2H6O' | 3.6 | 0.73 |

<sup>#</sup>Tentative identifications were assigned based on the highest NIST mass spectral library match with a match value (MV) ≥ 800; analytes with MV < 800 were reported as unknown.

mVOCs categorized in Category 1 were detected at significantly higher levels in the headspace of *P. aeruginosa* monoculture and co-culture compared to the media control and *A. fumigatus* monoculture, suggesting that they are likely produced by *P. aeruginosa*. A key metabolite in this Category, 4-methylquinazoline (Hit 57), is a known *P. aeruginosa* mVOC generated as a byproduct of 2-aminoacetophenone biosynthesis and has been identified as a potential breath biomarker for *P. aeruginosa* pulmonary infection.^52,53^ Sulfur- and nitrogen-containing compounds detected in this Category are also known *P. aeruginosa* metabolites. For example, dimethyl disulfide (Hit 16) and dimethyl trisulfide (Hit 69) are products of *P. aeruginosa* methionine metabolism.^2,54–59^ The nitrogen-containing compound, decanenitrile (Hit 15), is a known *P. aeruginosa* volatile reported to exert antifungal activity against mycelial growth and spore germination,^60^ suggesting its production may contribute to antagonistic interactions against *A. fumigatus*. Several Category 1 analytes had alternative NIST library matches with slightly lower MVs that correspond to compounds previously reported in the *P. aeruginosa* volatilome. For example, 2-aminoacetophenone was the second-highest library match (MV = 854) for Hit 2. This compound has been identified as a putative biomarker for *P. aeruginosa* and associated with chronic infection phenotypes.^53,53,54,61–64^ Similarly, the alternative assignment for Hit 34 (MV = 849), 2,3,5-trimethylpyridine, has previously been reported as a *P. aeruginosa* volatile metabolite.^54^ Although these alternative assignments are biologically plausible, library matching alone is insufficient for definitive compound identification, and additional structural confirmation is required.

Volatile metabolites in Category 2 were detected at significantly higher abundance in *A. fumigatus* monoculture compared to all of the other conditions, including the co-culture condition, indicating potential interkingdom mVOC interactions and cross-feeding. Among these metabolites, hydrocarbons such as hexadecane (Hit 94) can be consumed by *Pseudomonas spp*. as the sole carbon and energy source.^65^ The decreased abundance of hexadecane in the co-culture headspace relative to *A. fumigatus* monoculture, may indicate possible alkane vapor cross-feeding during co-cultivation. This conclusion is further supported by the presence of an alkane oxidation product, 1-hexadecanol (Hit 118), in the co-culture volatilome. Other hydrocarbons such as heptadecane (Hit 92) have been identified to be emitted by *Aspergillus flavus*, and known for its antimicrobial activity against pathogens.^65–67^ Two other notable metabolites in this Category, 4-ethoxybenzoic acid ethyl ester (Hit 41) and 2,6,10-trimethyl-tetradecane (Hit 129), have not been previously attributed to *A. fumigatus* or *P. aeruginosa* volatilomes, but both have been detected in metabolite extracts with reported antimicrobial activity against pathogens, including *P. aeruginosa*.^68,69^ Of the known *A. fumigatus* mVOCs, 1-octen-3-ol and isopentyl alcohol were not identified in the peak table because the peak table included only the 104 analytes that exhibited statistically significant differences in concentration across the four sample classes according to the experimental design. However, it is possible that they were among the 104 analytes but had NIST MVs below 800 and were therefore reported as unknowns.

The other three Categories represent metabolites consumed by *P. aeruginosa* only (Category 3), *A. fumigatus* only (Category 5), and both *P. aeruginosa* and *A. fumigatus* (Category 4). These mVOCs are chemically diverse, representing branched-chain aldehydes (Hits 8, 19, 26), aromatic aldehydes (Hits 4, 21, 27, 50), straight-chain aldehydes (Hits 1, 6, 7, 9, 10, 12-14, 33), alcohols (Hits 11, 20, 24, 25, 40, 61, 82, 103), alkanes (Hits 29, 43, 48), a ketone (Hit 58), a thioether (Hit 51), an aromatic ether (Hit 45), an aromatic quinone (Hit 116), and a carboxylic acid ester (Hit 60). Among these compounds, nonanal (Hit 14), benzaldehyde (Hit 21) and 3-methylbutanal (Hit 26) are known to be imported and metabolized by *P. aeruginosa*.^55,56^ Notably, aldehydes comprised a major portion of the compounds consumed in both monocultures and co-culture, indicating that aldehyde utilization may be a common and important feature of these organisms. The widespread consumption of aldehydes by both microbes in the co-culture conditions may suggest potential metabolic competition for aldehydes in nutrient-limited environments.

In addition to the five major Categories, the four analytes represented by gray open circles in the PCA score plot exhibited unique ternary codes but showed similar qualitative abundance patterns based on their PCA positions and signal profiles (Figure 7). Based on these similarities, the four gray open-circle analytes were further grouped into an additional Category 6, characterized by significantly higher abundance in both monocultures and co-culture compared to the media control, representing a related biological pattern not captured by the five recurring ternary-code Categories. Among these analytes, 2-nonanone (Hit 66) exhibited a distinct abundance pattern with high levels in all three microbial cultures relative to media, which is consistent with previously reports of this molecule as a signature volatile metabolite of both *P. aeruginosa* and *A. fumigatus*.^57,70,71^ Notably, three major abundance patterns were not observed in our analysis: (1) metabolites significantly higher in *P. aeruginosa* compared to all other conditions, (2) metabolites produced in *A. fumigatus* monoculture and co-culture, and (3) volatiles uniquely enriched under co-culture conditions. The absence of the first two patterns suggests that microbial cross-feeding may be one-sided, where metabolites produced by *A. fumigatus* are consumed by *P. aeruginosa* (Category 2) but not the reverse. Meanwhile, the absence of a diagnostic co-culture pattern indicates that the tested volatilome conditions and analysis method is limited in identifying potential volatile biomarkers of *P. aeruginosa* and *A. fumigatus* co-infection.Together, these findings support the role of mVOCs as mediators of interkingdom interactions. The observed patterns of production, depletion, and culture-specific abundance suggest that these mVOCs contribute to microbial competition, resource acquisition, and metabolic cross-feeding. In particular, volatiles that can serve as carbon and energy sources are important for microbial survival in hostile environments like the lungs. *P. aeruginosa* and *A. fumigatus* are often co-present in the airways of individuals with cystic fibrosis (CF),^72–74^ where competition with each other for nutritional resources is critical for microbial survival and colonization. The potential for volatile metabolite cross-feeding, where metabolites produced by *A. fumigatus* were consumed by *P. aeruginosa*, together with the lack of detectable growth inhibition in the co-culture conditions (Figure S5), suggest that mVOC-mediated interkingdom interactions may support the fitness of these microbes in nutrient-limited niches. Importantly, the biological effects of these mVOC signals and interactions are likely dependent on the environmental context. For instance, some of these volatiles that are toxic or inhibitory to either microbe in a nutrient deficient environment, such as the lung, may not exert similar effects in the nutrient-rich experimental conditions used in this study.

## CONCLUSION

We developed versatile microbial co-culture devices and a systematic data analysis workflow for investigating *P. aeruginosa* and *A. fumigatus* monoculture and co-culture volatilomes. Our double-layer co-culture device enables isolation and characterization of mVOC-mediated interactions between microbes at distance, preventing spore-derived physical contacts. Exogenously supplementing known *A. fumigatus* VOCs, 1-octen-3-ol and isopentyl alcohol, inhibited the growth and biofilm formation of *P. aeruginosa*. Although no measurable physiological changes in *P. aeruginosa* growth and biofilm formation were detected during co-cultivation with *A. fumigatus* under the tested conditions, our device enabled efficient, high-throughput sampling and characterization of their volatilomes through direct compatibility with GC autosamplers. Using untargeted tile-based F-ratio analysis on the GC×GC-TOFMS data, we identified 198 volatiles which differed across at least two of the four sample classes (M, PA, AF, and CC); 104 of which met the significance threshold (*p*-value < 0.01). Our systematic probability-based ternary-code approach further classified these 104 volatiles into five major Categories with distinct abundance patterns, enabling interpretation of differences across microbial volatilomes and sample conditions. The resulting volatile profiles were consistent with previously reported *P. aeruginosa* and *A. fumigatus* volatilomes, while our approach expanded the known volatilomes and provided insights into potential roles of individual mVOCs in microbial interactions. Overall, these microbial co-culture devices and analytical workflow provide a versatile approach for investigating mVOC exchanges across diverse microbial systems. This platform may facilitate future studies on the roles of volatile metabolites in complex communities and future efforts to identify volatile biomarkers for non-invasive diagnosis of microbial infections, while our analytical approach provides a framework for classifying multidimensional datasets across diverse omics applications and sample classes.

## ASSOCIATED CONTENT

### SUPPORTING INFORMATION

The Supporting Information (pdf) is available free of charge on the ACS Publications website.

Supporting Information: detailed device dimensions and assembly; validation of the device; co-culture growth and biofilm assessments; Monte Carlo simulation results; PCA results; chromatographic profiles and signal intensities for the 104 ANOVA-significant analytes; ternary-code patterns summarized in tables; and additional supporting data (PDF). Supporting files: Device CAD files (.f3d); PTFE membrane design files (PDF); Raw GC×GC-TOFMS data for all six biological replicates of each sample class are available through Zenodo at 10.5281/zenodo.23581553.

## AUTHOR INFORMATION

### Author Contributions

The manuscript was written through contributions of all authors, and all authors have given approval to the final version of the manuscript. TW: Writing – original draft review & editing, Visualization, Validation, Software, Methodology, Investigation, Formal analysis, Data curation, Conceptualization. WM: Writing – original draft review & editing, Visualization, Validation, Software, Methodology, Investigation, Formal analysis, Data curation, Conceptualization. JMW: Writing − review and editing, Formal analysis, Investigation and Conceptualization. JCT: Writing – review & editing, Methodology, Investigation and Conceptualization. HC: Investigation, Data curation. JC: Investigation, Data curation. KYU: Investigation, Data curation. UNL: Investigation and Conceptualization. BNS: Writing – review & editing, Advising, Conceptualization. NPK: Writing – review & editing, Advising, Conceptualization. EB: Advising, Methodology, Conceptualization. RES: Writing – review & editing, Supervision, Resources, Project administration, Methodology, Investigation, Funding acquisition, Conceptualization. ABT: Writing – review & editing, Supervision, Resources, Project administration, Methodology, Investigation, Funding acquisition, Conceptualization. LJR: Writing – review & editing, Supervision, Resources, Project administration, Methodology, Investigation, Funding acquisition, Conceptualization.

## ACKNOWLEDGEMENT

We want to acknowledge the M. J. Murdock Diagnostic Foundry for Translational Research. This publication was supported by the National Institutes of Health (NIH) through R35GM128648 (A.B.T.) and R35GM150913 (L.J.R.). The content is solely the responsibility of the authors and does not necessarily represent the official views of the National Institutes of Health or other funders.

