## Supporting information for "Development of an Enclosed Microbial Co-culture Device and GC×GC-TOFMS Workflow to Investigate Volatile Organic Compound Exchange"

**Table of Contents**

**Figure S1.** Schematic diagrams of the microbial co-culture devices S3

**Figure S2.** Zoomed-in views of the microbial co-culture device schematics shown in Figure S1 S4

**Figure S3.** Validation of double-layer co-culture device for preventing spore migration. S5

**Figure S4.** VOCs diffuse across the membrane and induce physiological changes of *P. aeruginosa* 14. S6

**Figure S5.** Growth of *P. aeruginosa* was not affected upon 24h exposure to mVOCs produced by *A. fumigatus*. S7

**Figure S6.** Distribution of Log10(*p-value*) from one-way ANOVA at the 99% confidence level plotted against hit index. S8

**Figure S7.** Hits from Category 1 and its ternary-pattern subgroups. S9

**Figure S8.** Hits from Category 2 and its ternary-pattern subgroups. S10-11

**Figure S9.** Hits from Category 3 and its ternary-pattern subgroups. S12-13

**Figure S10.** Hits from Category 4 and its ternary-pattern subgroups. S14-15

**Figure S11.** Hits from Category 5 and its ternary-pattern subgroups. S16

**Figure S12.** Sixteen hits with unique ternary patterns. S17-18

Theoretical distribution and Monte Carlo simulation of ternary-code recurrence. S19

**Figure S13.** Monte Carlo simulation of ternary-code pattern recurrence under random assignment. S20

**Figure S14.** PCA of the 104 hits meeting the p-value < 0.01 criterion in one-way ANOVA. S21

**Table 1.** Ternary-code patterns observed among the 94 hits with p-value ≥ 0.01. S22

**Table 2.** Ternary-code patterns observed among the 104 hits with p-value < 0.01. S23

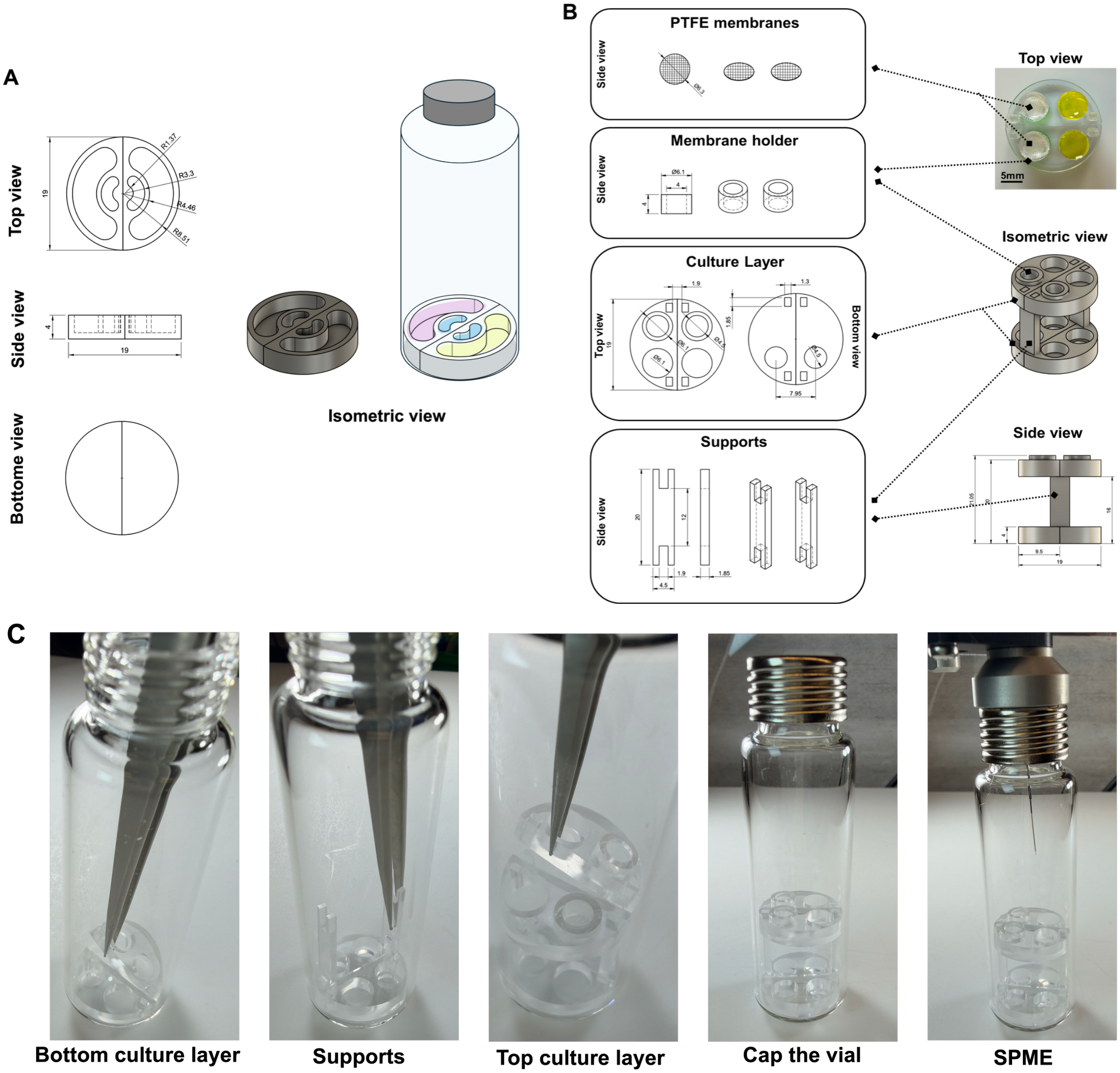


**Figure S1.** Schematic diagrams of microbial co-culture devices. Schematic diagrams illustrating the dimensions of the single-layer (A) and the double-layer (B) microbial coculture devices, shown from top, side, bottom and isometric views. (C) Photographs showing the assembly process of the double-layer device. All dimensions are in mm. PTFE: polytetrafluoroethylene; SPME: solid-phase microextraction.

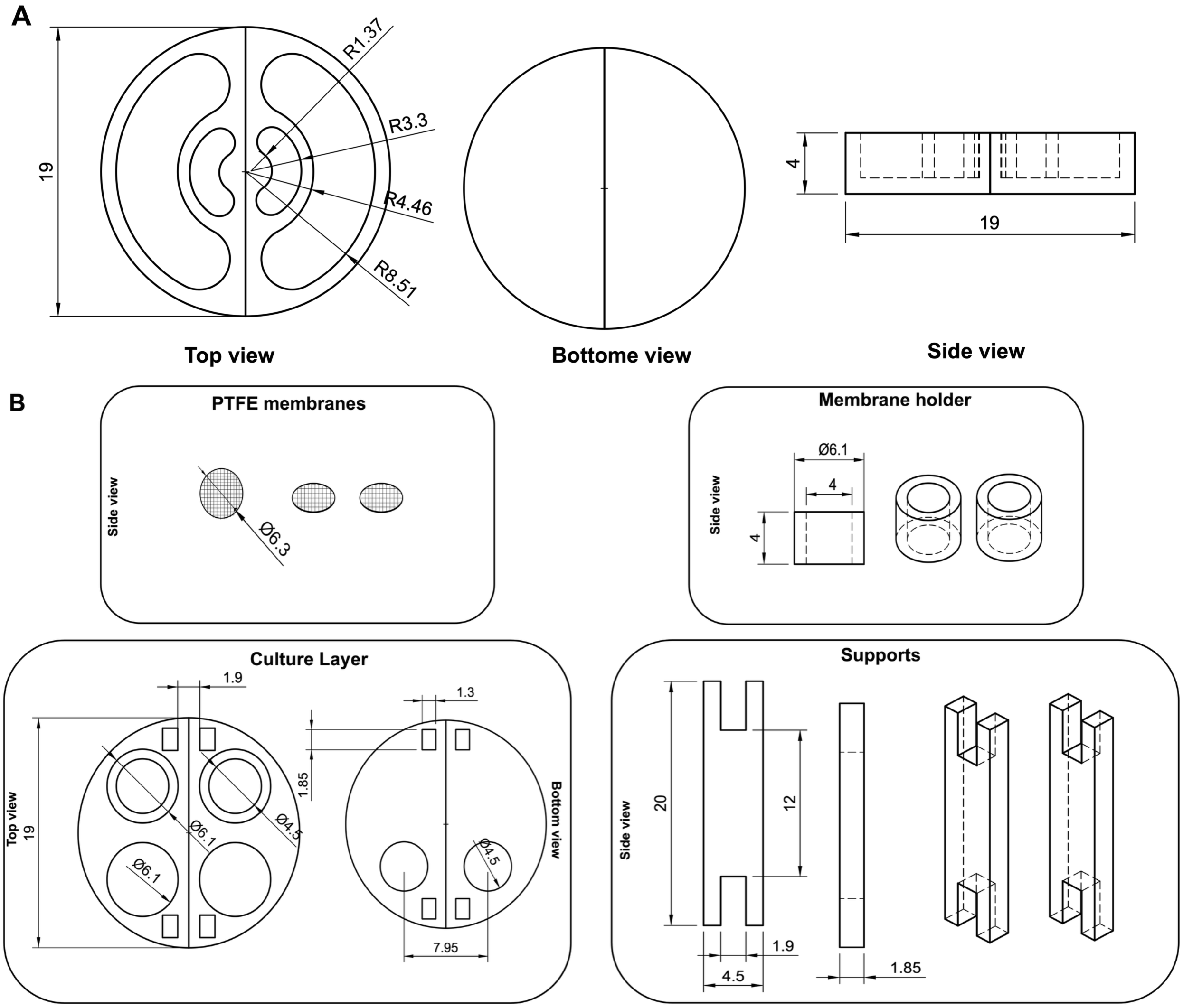


**Figure S2.** Zoomed-in views of the microbial co-culture device schematics shown in Figure S1. Schematic diagrams illustrating the dimensions of the single-layer (A) and the double-layer (B) microbial co-culture devices. All dimensions are in mm. PTFE: polytetrafluoroethylene.

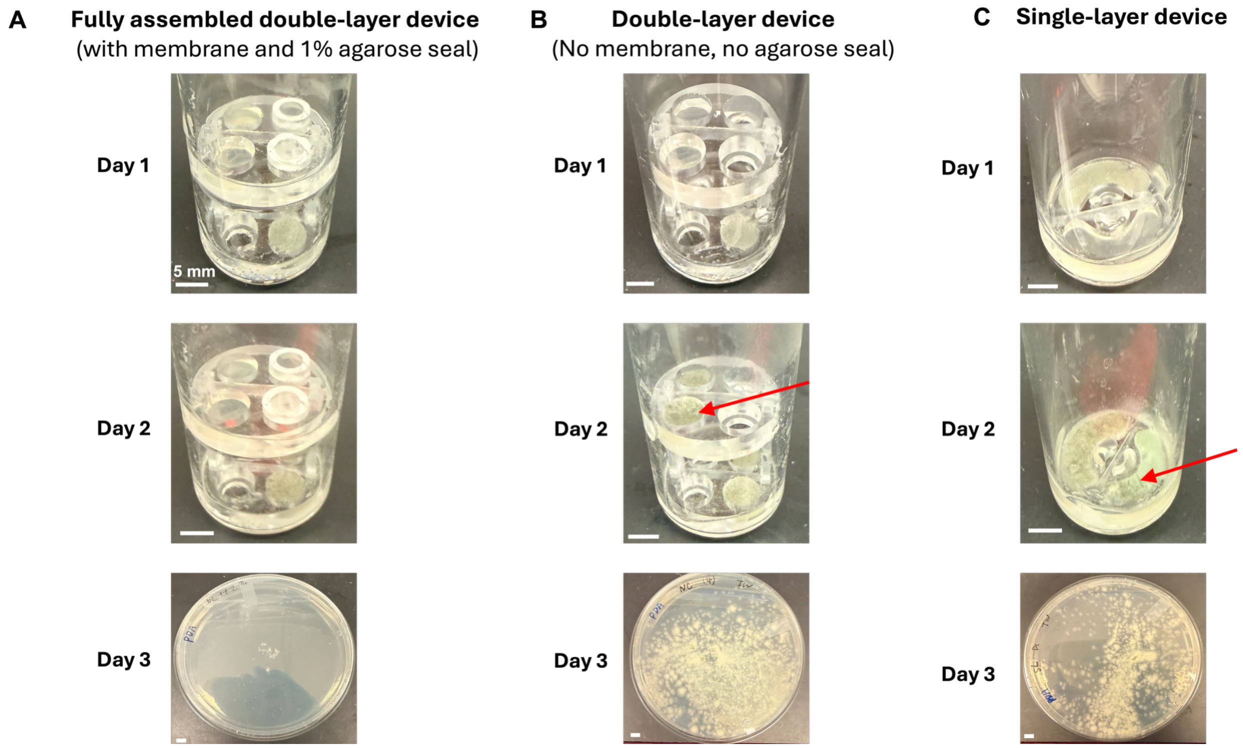


**Figure S3.** Validation of double-layer co-culture device for preventing spore migration. Photographs of the fully assembled double-layer co-culture device (A), double-layer co-culture device without membrane and agarose seal (B) and single-layer co-culture device (C) on day 1 and day 2 after inoculation of *A. fumigatus* 293 into the bottom culture wells of the double-layer devices and one culture well of the single layer device; Uninoculated PDA in the top culture wells (A, B) and the neighboring culture well (C) served as a spore-capture medium to monitor spore migration. On day 2, the spore-captured media were transferred onto a Petri-dish containing sterile PDA and incubated at 37 °C for 24 hours to allow any contaminating spore to grow. Photographs of the Petri-dishes were taken on day 3. Samples from (A) showing no visible fungal growth were washed with 0.1 v/v tween 80 and inspected microscopically to confirm the absence of fungal growth. Red arrows indicate spore contamination. All the systems were incubated at 37 °C with PDA used as a growth medium. White scale bars indicate 5 mm.

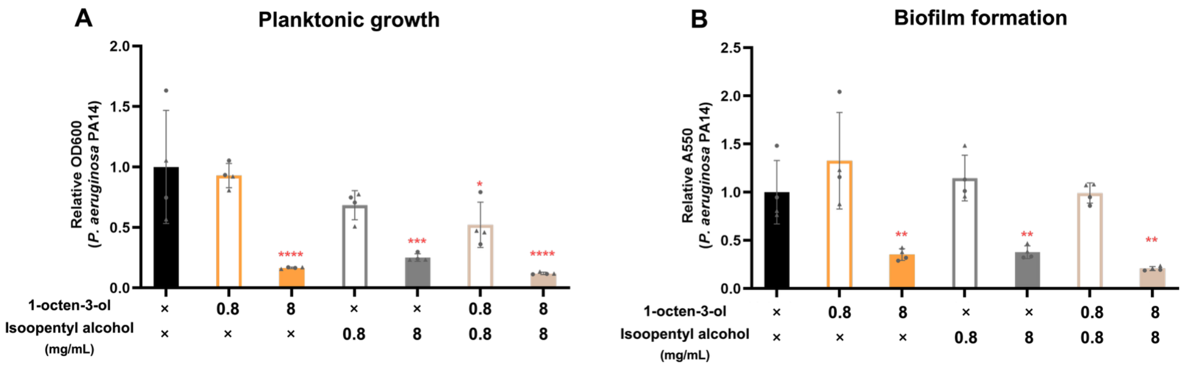


**Figure S4.** Diffusion of VOCs across the membrane and induce physiological changes of *P. aeruginosa* PA14. (A) Schematics of mVOC-addition experimental conditions. (B) Planktonic growth of *P. aeruginosa* PA14, after 24 h exposure to two different doses (0.8 mg/mL and 8 mg/mL) of 1-octen-3-ol, isopentyl alcohol, and their equal mixture. Planktonic growth was assessed and quantified by the planktonic cultures at 600 nm. (C) Biofilm formation of *P. aeruginosa* PA14 after 24 h exposure to two different doses (0.8 mg/mL and 8 mg/mL) of 1-octen-3-ol, isopentyl alcohol. Mean values were calculated for four samples originating from two culture wells in two independent devices; data points with the same shape indicate samples from the same device. The mean values were normalized to the vehicle control (black bars). The error bars represent the standard error of the mean. Statistical analysis was performed comparing each condition to the vehicle control using one-way ANOVA with Tukey's multiple comparisons test. Statistical differences are shown with red asterisks (no asterisk, $p$-value > 0.05; *, $p$-value < 0.05; **, $p$-value < 0.01; ***, $p$-value < 0.001; ****, $p$-value < 0.0001).

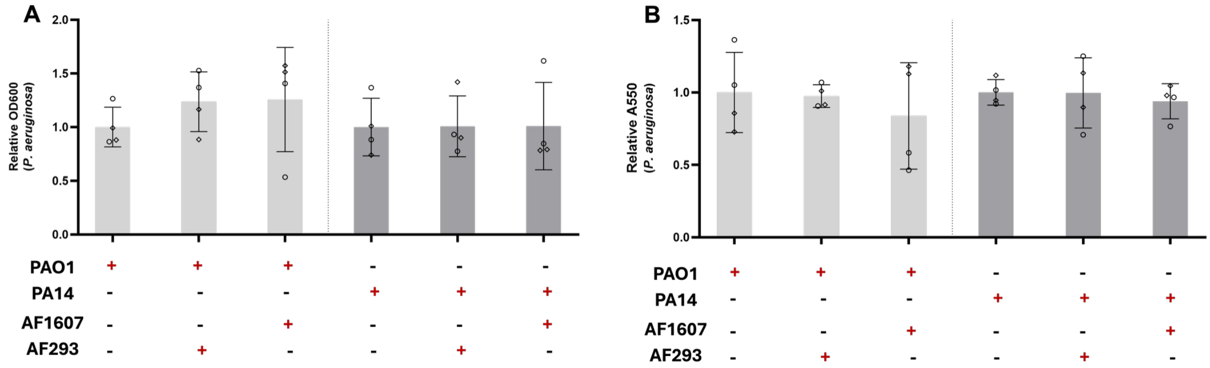


**Figure S5.** Growth of *P. aeruginosa* was not affected upon 24 h exposure to mVOCs produced by *A. fumigatus*. (A) Planktonic growth of *P. aeruginosa* 24 h after co-culture with *A. fumigatus*. (B) Biofilm formation of *P. aeruginosa* 24 h after co-culture with *A. fumigatus*. Mean values were calculated for four samples originating from two culture wells in two independent devices; data points with the same shape indicate samples from the same device. The presence (+) or absence (-) of each indicated microorganism in the device is shown below the corresponding bars. The mean values were normalized to corresponding *P. aeruginosa* monocultures. The error bars represent standard error of the mean. Statistical analysis was performed using one-way ANOVA with Tukey's multiple comparisons test (no asterisk, p-value > 0.05).

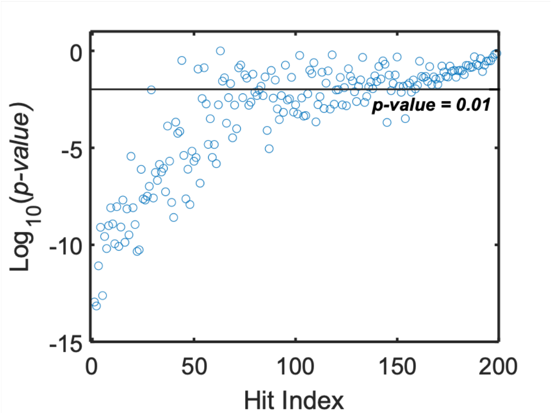


| *p-value* | Number of Hits | Accumulated Number of Hits |
|---|---|---|
| p >= 0.05 | 53 | 198 |
| 0.01 <= p < 0.05 | 41 | 145 |
| 0.001 <= p < 0.01 | 35 | **104** |
| 0.0001 <= p < 0.001 | 13 | 69 |
| p < 0.0001 | 56 | 56 |

**Figure S6.** Distribution of $\log_{10}$(*p*-value) from one-way ANOVA at the 99% confidence level plotted against hit index. The horizontal line indicates the *p*-value = 0.01 significance threshold. Of the 198 hits evaluated, 104 met the *p*-value < 0.01 criterion and were carried forward for pairwise t-test analysis. The table on the right summarizes the number of hits within each *p*-value range and the corresponding cumulative counts.

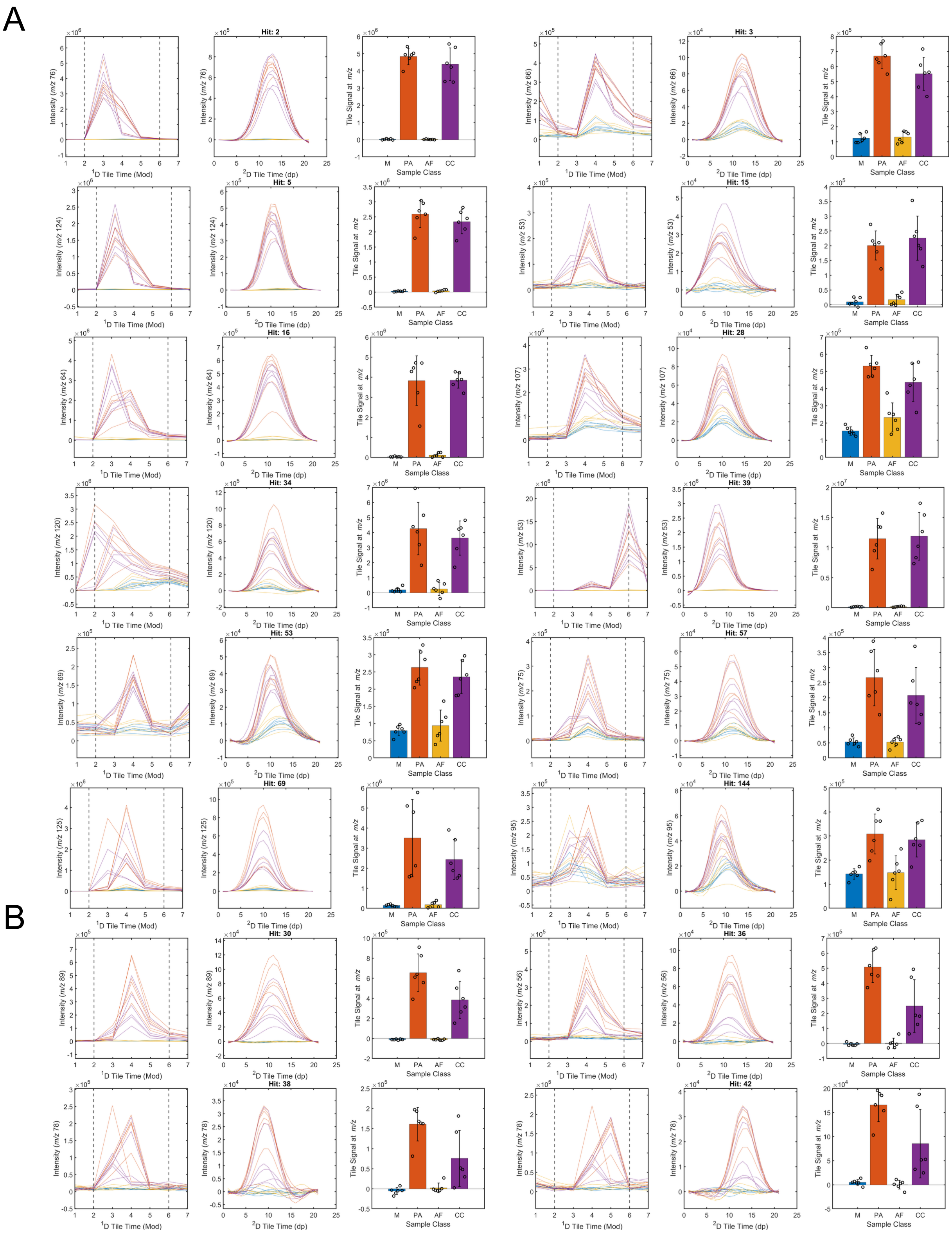


**Figure S7.** Hits from Category 1 and its ternary-pattern subgroups. (A) Group A (n = 12). (B) Group H (n = 4). For both groups, each hit is presented using three panels showing the $^{1}$D and $^{2}$D tile chromatographic profiles and the tile signal intensity across the sample classes (M, PA, AF, and CC).

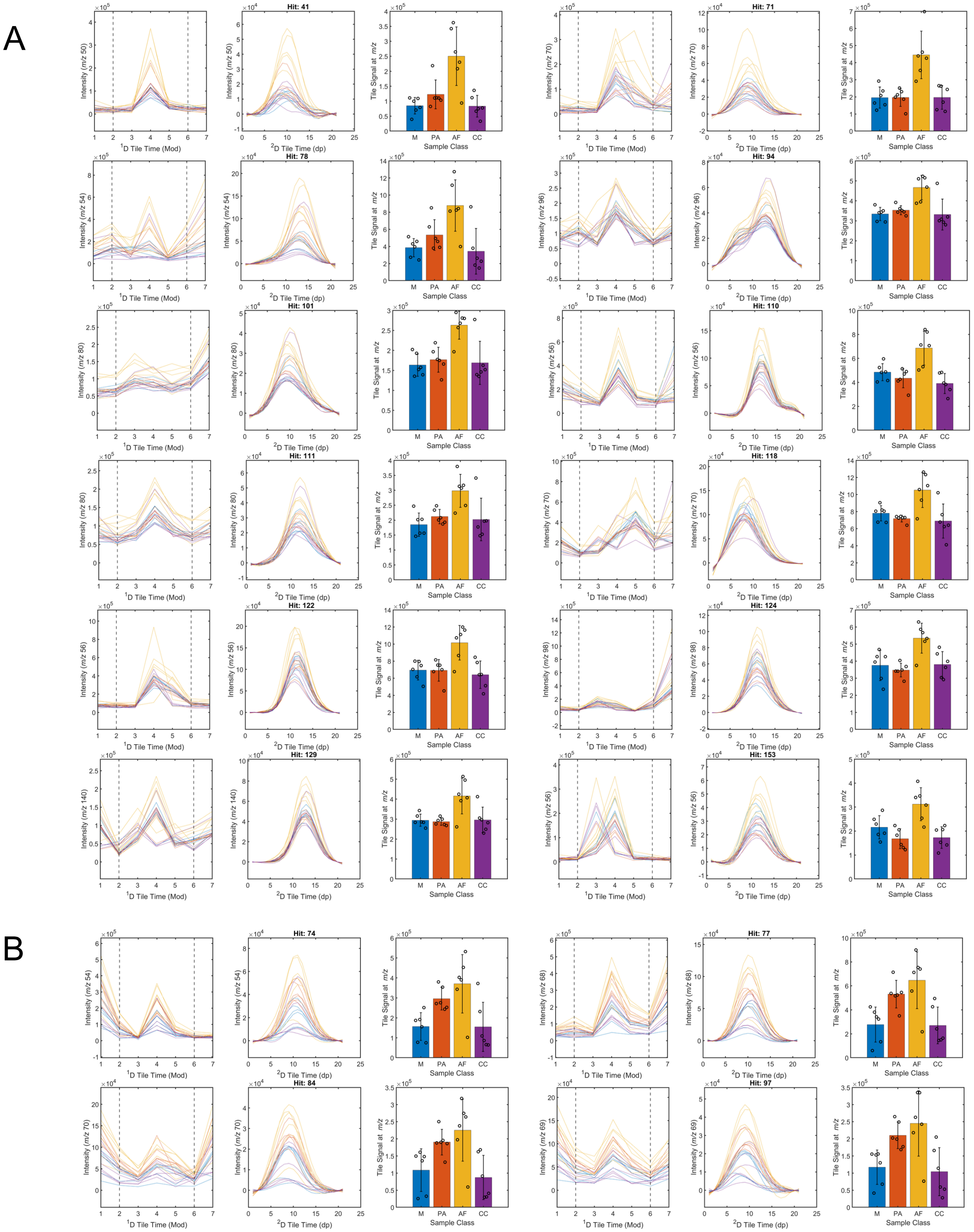
A
B
Hit: 41
Hit: 71
Hit: 78
Hit: 94
Hit: 101
Hit: 110
Hit: 111
Hit: 118
Hit: 122
Hit: 124
Hit: 129
Hit: 153
Hit: 74
Hit: 77
Hit: 84
Hit: 97
Intensity (m/z 50)
Intensity (m/z 70)
Intensity (m/z 54)
Intensity (m/z 96)
Intensity (m/z 80)
Intensity (m/z 56)
Intensity (m/z 98)
Intensity (m/z 140)
Intensity (m/z 68)
Intensity (m/z 69)
Tile Signal at m/z
1D Tile Time (Mod)
2D Tile Time (dp)
Sample Class
M
PA
AF
CC

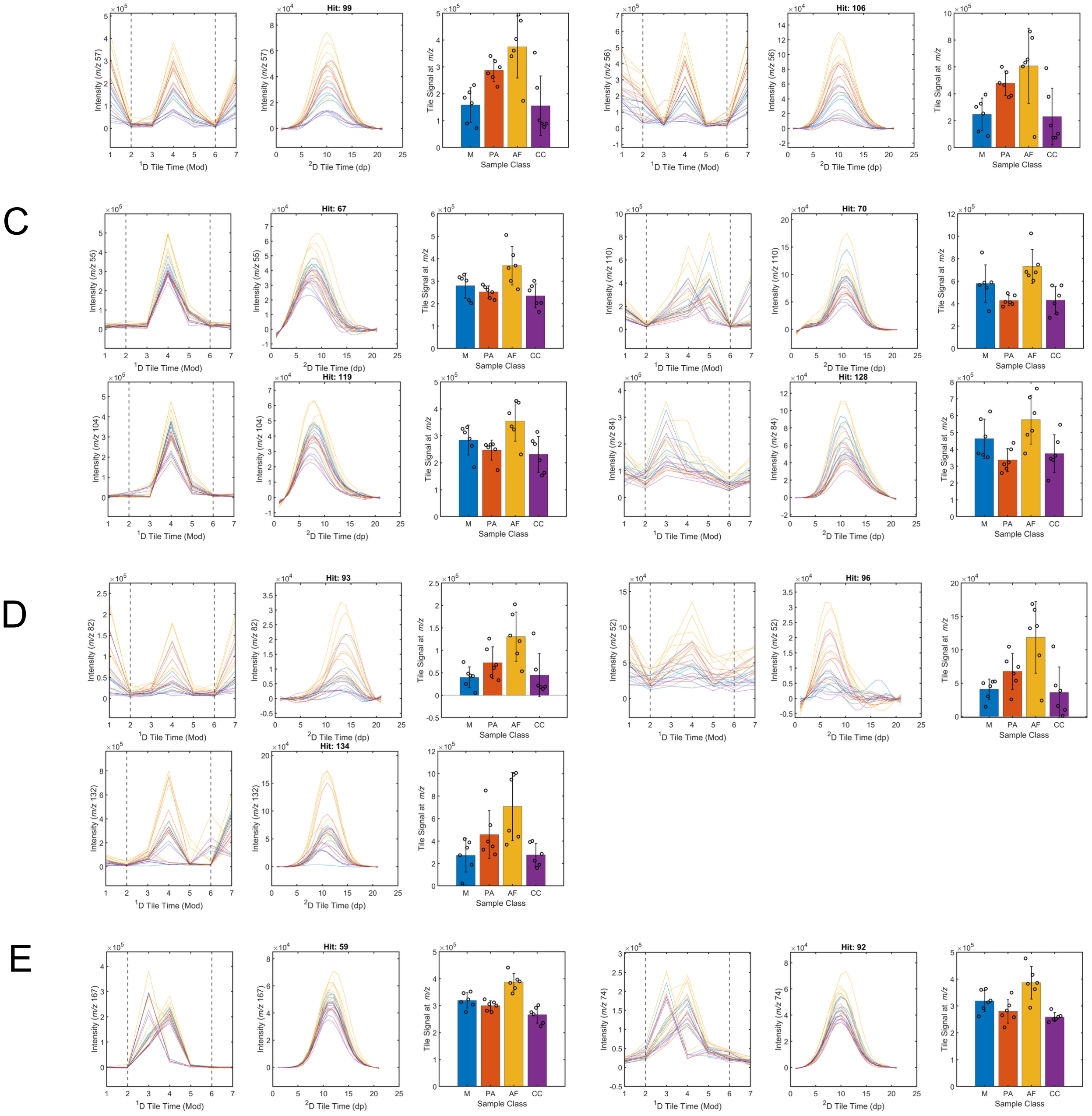


**Figure S8.** Hits from Category 2 and its ternary-pattern subgroups. (A) Group B (n = 12). (B) Group F (n = 6). (C) Group I (n = 4). (D) Group K (n = 3). (E) Group M (n = 2). For all five groups, t each hit is presented using three panels showing the $^{1}$D and $^{2}$D tile chromatographic profiles and the tile signal intensity across the sample classes (M, PA, AF, and CC).

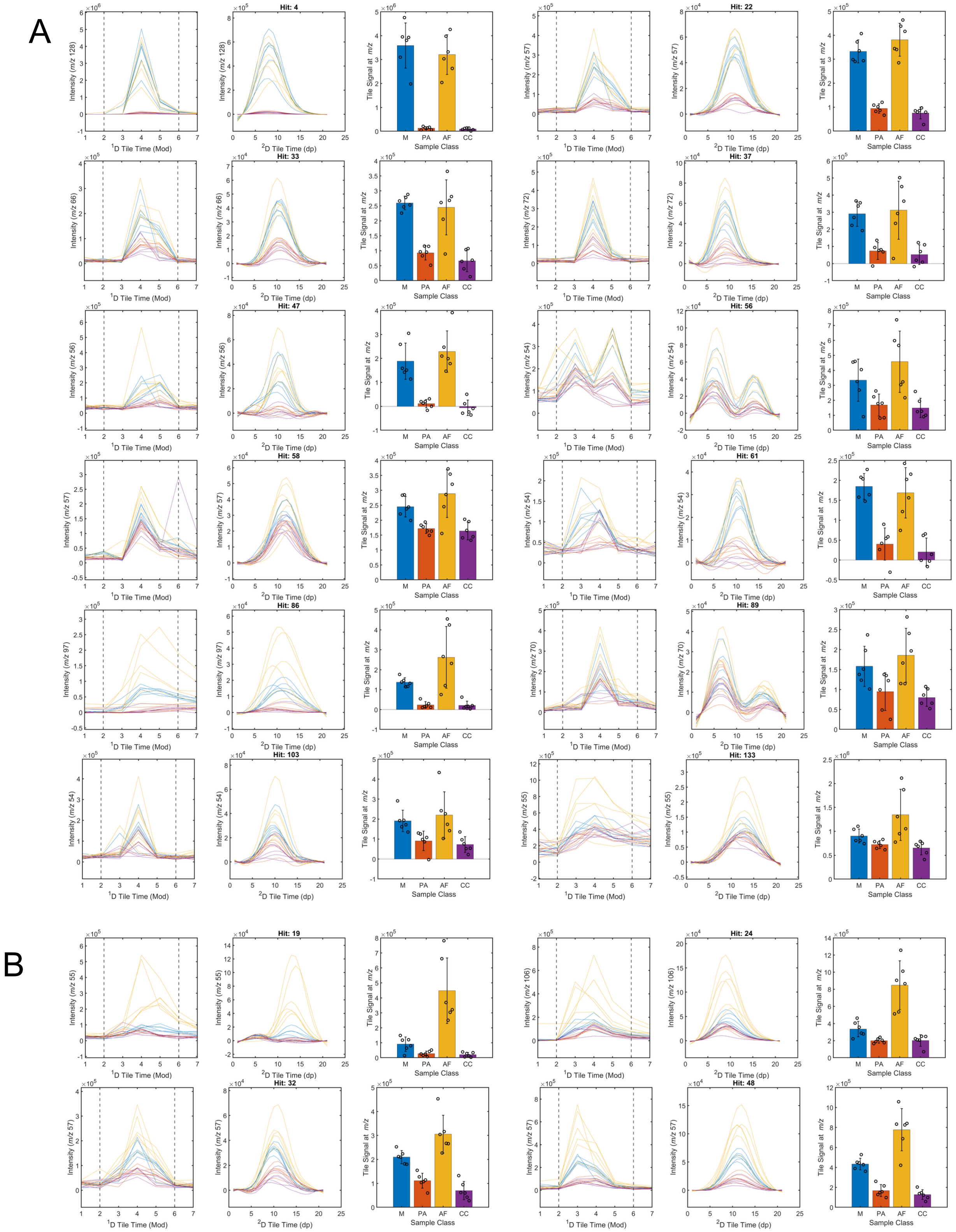
A
Hit: 4
Hit: 22
Hit: 33
Hit: 37
Hit: 47
Hit: 56
Hit: 58
Hit: 61
Hit: 86
Hit: 89
Hit: 103
Hit: 133
B
Hit: 19
Hit: 24
Hit: 32
Hit: 48
Intensity (m/z 128)
Intensity (m/z 57)
Intensity (m/z 66)
Intensity (m/z 72)
Intensity (m/z 56)
Intensity (m/z 54)
Intensity (m/z 97)
Intensity (m/z 70)
Intensity (m/z 55)
Intensity (m/z 106)
Tile Signal at m/z
¹D Tile Time (Mod)
²D Tile Time (dp)
Sample Class
M
PA
AF
CC

C

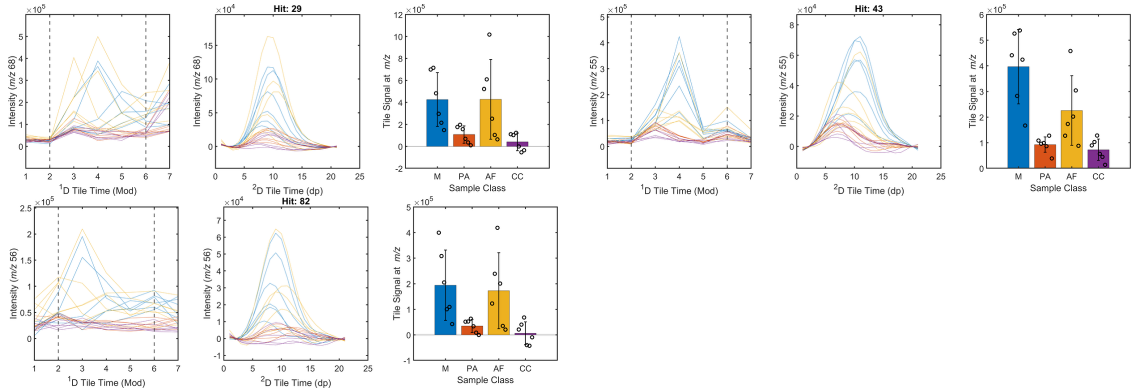


**Figure S9.** Hits from Category 3 and its ternary-pattern subgroups. (A) Group C (n = 12). (B) Group J (n = 4). (C) Group L (n = 3). For all three groups, each hit is presented using three panels showing the $^{1}$D and $^{2}$D tile chromatographic profiles and the tile signal intensity across the sample classes (M, PA, AF, and CC).

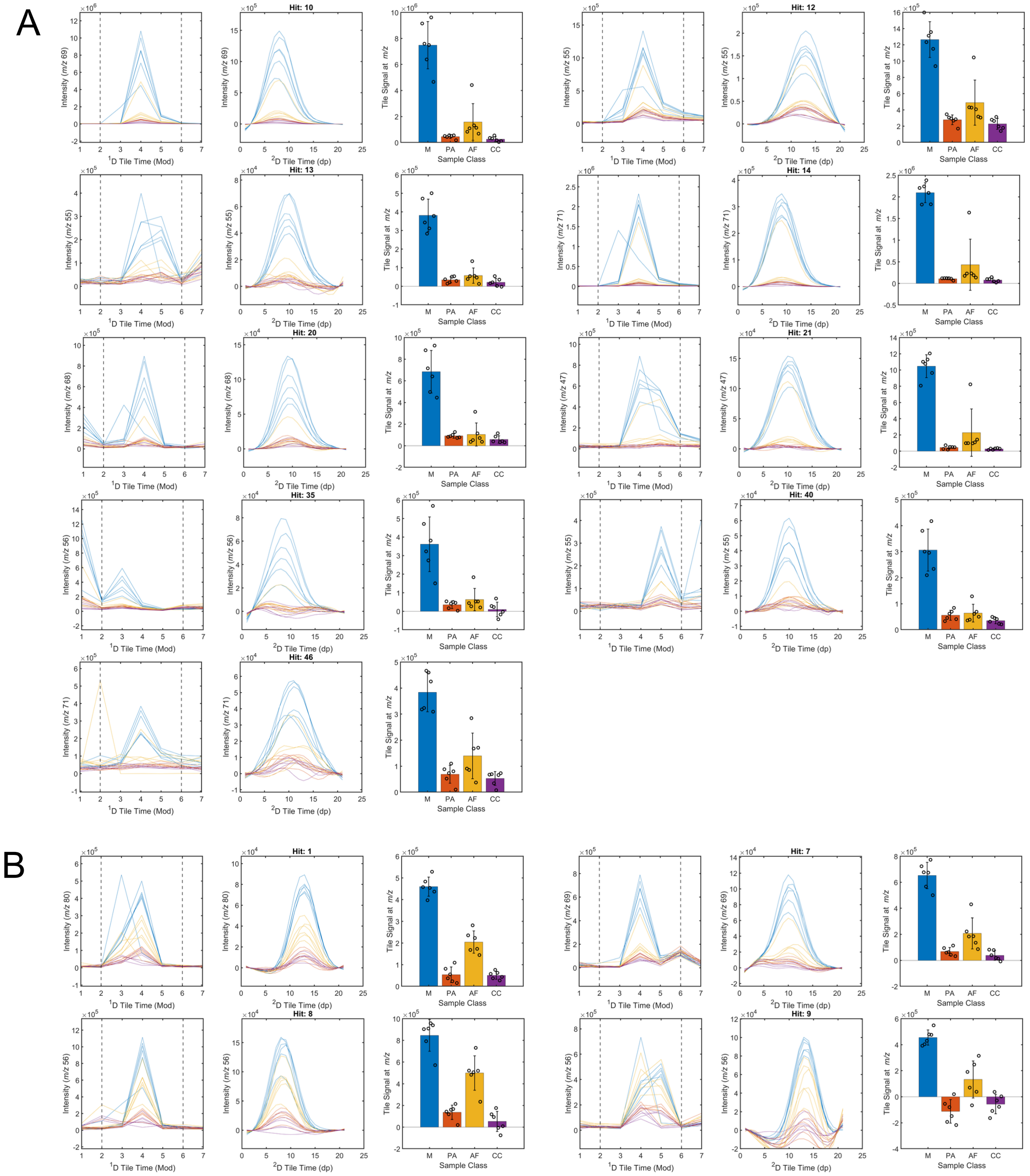
A
Hit: 10
Hit: 12
Hit: 13
Hit: 14
Hit: 20
Hit: 21
Hit: 35
Hit: 40
Hit: 46
B
Hit: 1
Hit: 7
Hit: 8
Hit: 9
Intensity (m/z 69)
Intensity (m/z 55)
Intensity (m/z 71)
Intensity (m/z 68)
Intensity (m/z 47)
Intensity (m/z 56)
Intensity (m/z 80)
Tile Signal at m/z
¹D Tile Time (Mod)
²D Tile Time (dp)
Sample Class
M
PA
AF
CC

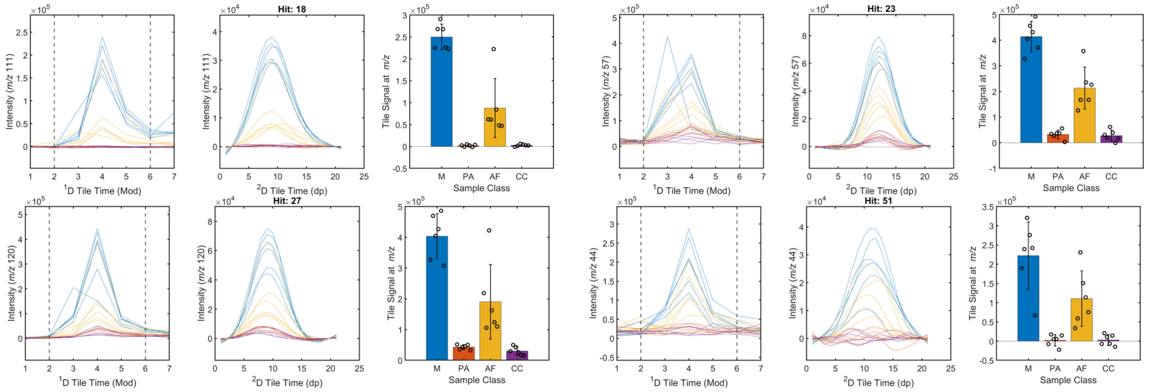


C

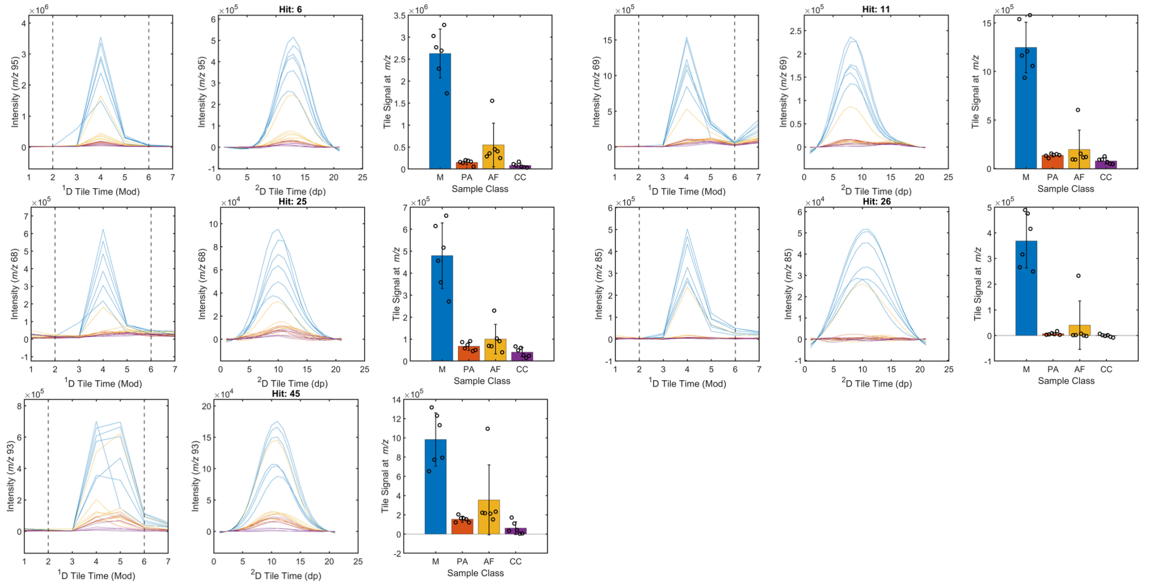


D

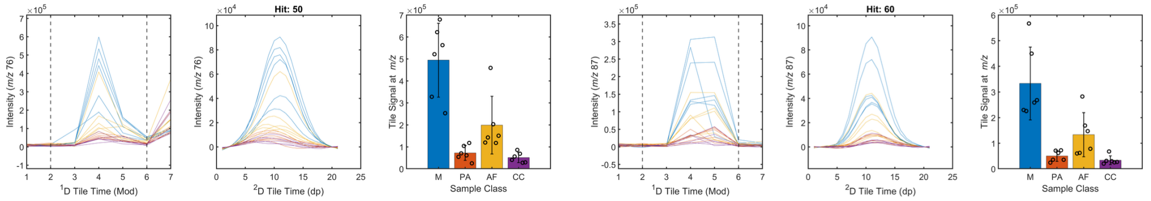


**Figure S10.** Hits from Category 4 and its ternary-pattern subgroups. (A) Group D (n = 9). (B) Group E (n = 8). (C) Group G (n = 5). (D) Group O (n = 2). For all four groups, each hit is presented using three panels showing the $^1$D and $^2$D tile chromatographic profiles and the tile signal intensity across the sample classes (M, PA, AF, and CC).

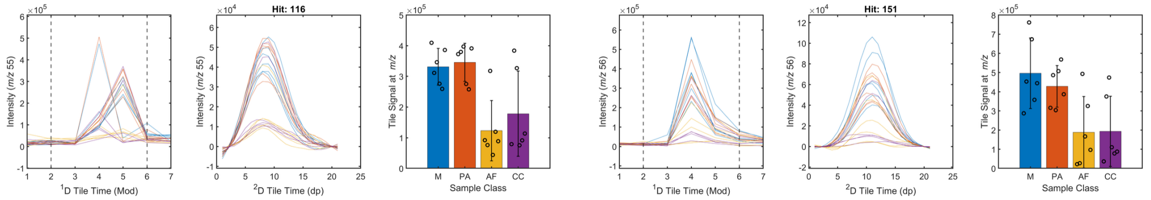


**Figure S11.** Hits from Category 5 and its ternary-pattern subgroup Group N (n = 2). Each hit is presented using three panels showing the $^{1}$D and $^{2}$D tile chromatographic profiles and the tile signal intensity across the sample classes (M, PA, AF, and CC).

| M vs. PA | M vs. AF | M vs. CC | PA vs. AF | PA vs. CC | AF vs. CC |
|---|---|---|---|---|---|
| 1 | 0 | 1 | -1 | 1 | 1 |

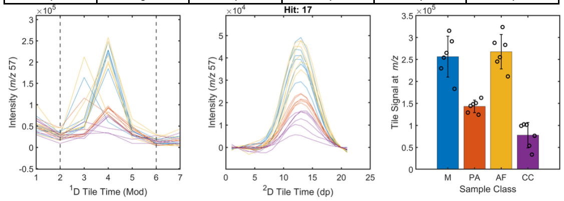


| M vs. PA | M vs. AF | M vs. CC | PA vs. AF | PA vs. CC | AF vs. CC |
|---|---|---|---|---|---|
| 1 | -1 | 1 | -1 | 1 | 1 |

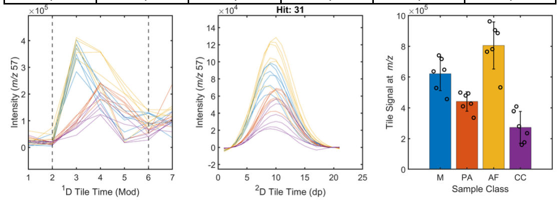


| M vs. PA | M vs. AF | M vs. CC | PA vs. AF | PA vs. CC | AF vs. CC |
|---|---|---|---|---|---|
| 1 | 1 | 1 | 0 | 1 | 1 |

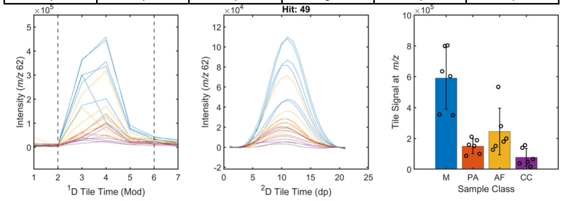


| M vs. PA | M vs. AF | M vs. CC | PA vs. AF | PA vs. CC | AF vs. CC |
|---|---|---|---|---|---|
| 1 | 0 | 0 | -1 | 0 | 1 |

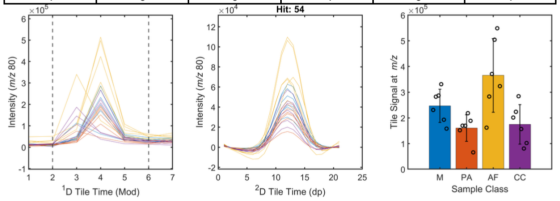


| M vs. PA | M vs. AF | M vs. CC | PA vs. AF | PA vs. CC | AF vs. CC |
|---|---|---|---|---|---|
| -1 | -1 | 0 | 0 | 0 | 0 |

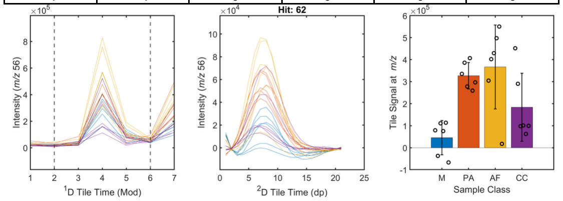


| M vs. PA | M vs. AF | M vs. CC | PA vs. AF | PA vs. CC | AF vs. CC |
|---|---|---|---|---|---|
| -1 | -1 | -1 | 0 | 0 | 0 |

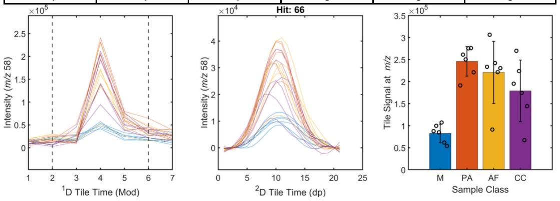


| M vs. PA | M vs. AF | M vs. CC | PA vs. AF | PA vs. CC | AF vs. CC |
|---|---|---|---|---|---|
| 0 | 0 | 1 | 0 | 0 | 1 |

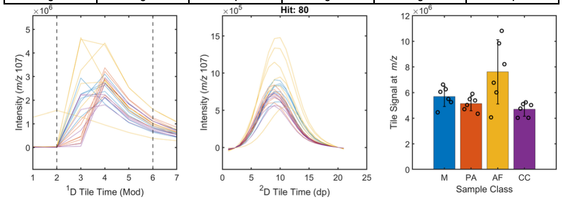


| M vs. PA | M vs. AF | M vs. CC | PA vs. AF | PA vs. CC | AF vs. CC |
|---|---|---|---|---|---|
| --1 | -1 | -1 | 0 | 1 | 0 |

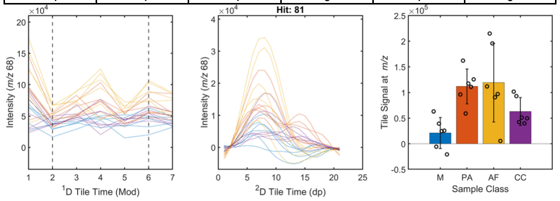


| M vs. PA | M vs. AF | M vs. CC | PA vs. AF | PA vs. CC | AF vs. CC |
|---|---|---|---|---|---|
| -1 | -1 | -1 | 1 | 1 | 0 |

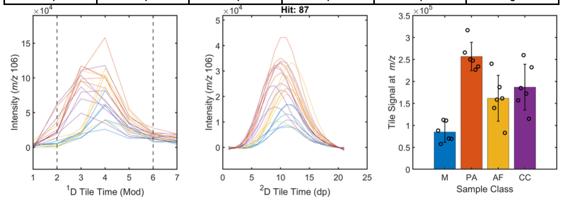


| M vs. PA | M vs. AF | M vs. CC | PA vs. AF | PA vs. CC | AF vs. CC |
|---|---|---|---|---|---|
| -1 | 0 | -1 | 1 | 0 | 0 |

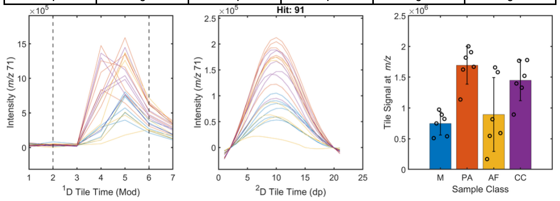


| M vs. PA | M vs. AF | M vs. CC | PA vs. AF | PA vs. CC | AF vs. CC |
|---|---|---|---|---|---|
| -1 | -1 | -1 | 0 | -1 | 0 |

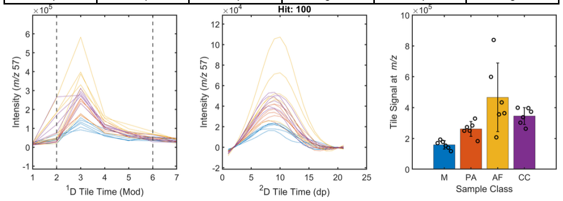


| M vs. PA | M vs. AF | M vs. CC | PA vs. AF | PA vs. CC | AF vs. CC |
|---|---|---|---|---|---|
| -1 | 0 | 0 | 1 | 1 | 0 |

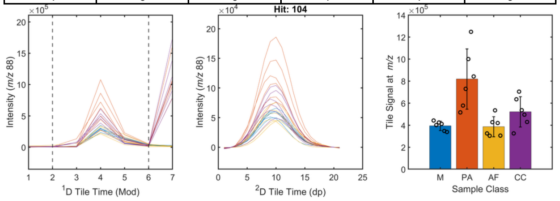


| M vs. PA | M vs. AF | M vs. CC | PA vs. AF | PA vs. CC | AF vs. CC |
|---|---|---|---|---|---|
| -1 | -1 | -1 | 0 | 0 | 1 |

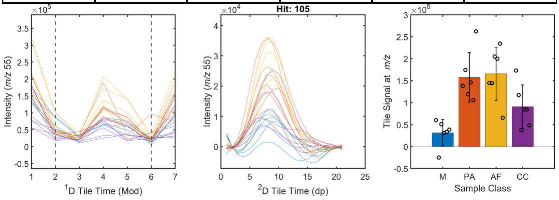


| M vs. PA | M vs. AF | M vs. CC | PA vs. AF | PA vs. CC | AF vs. CC |
|---|---|---|---|---|---|
| 1 | -1 | 0 | -1 | 0 | 1 |

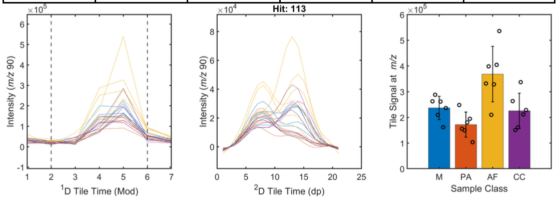

| M vs. PA | M vs. AF | M vs. CC | PA vs. AF | PA vs. CC | AF vs. CC |
| --- | --- | --- | --- | --- | --- |
| 0 | 0 | 0 | -1 | 0 | 0 |

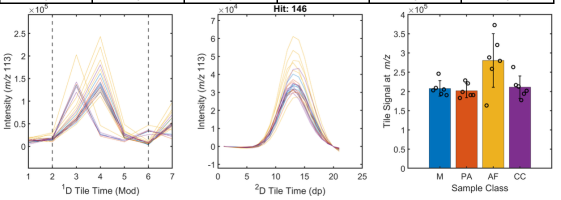


| M vs. PA | M vs. AF | M vs. CC | PA vs. AF | PA vs. CC | AF vs. CC |
| --- | --- | --- | --- | --- | --- |
| 1 | 0 | 1 | 0 | 0 | 0 |

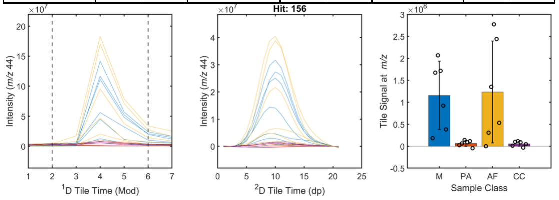


**Figure S12.** Sixteen hits with unique ternary patterns. For each hit, the six-digit ternary code (top) and three panels show representative ¹D and ²D tile chromatographic profiles and the tile signal intensity across sample classes (M, PA, AF, CC).

**Theoretical distribution and Monte Carlo simulation of ternary-code recurrence.** To evaluate the expected recurrence of ternary-code patterns under random assignment, 104 analytes were independently assigned with equal probability to the 729 possible six-digit ternary codes.[1,2] For a specific ternary code, the number of assigned analytes, X, follows a binomial distribution.[1,2] The probability that a specific code occurs exactly $k$ times is

$$P(X=k) = \binom{104}{k}\left(\frac{1}{729}\right)^{k}\left(\frac{728}{729}\right)^{104-k}$$

Because there are 729 possible ternary codes, the expected number of codes occurring exactly $k$ times is

$$E_k = 729P(X=k) = 729\binom{104}{k}\left(\frac{1}{729}\right)^{k}\left(\frac{728}{729}\right)^{104-k}$$

For example, for $k = 2$, $P(X = 2) = 0.00876$, corresponding to an expected 6.39 ternary codes occurring exactly twice per simulation.

To compare these theoretical expectations with simulation, the random-assignment procedure was repeated $10^6$ times. Within each simulation, only distinct code patterns occurring at least once were counted, and each observed pattern was classified according to its number of occurrences among the 104 analytes. For example, if 102 analytes were assigned unique codes and the remaining two analytes shared the same code, the simulation would contain 103 distinct observed code patterns: 102 occurring once and one occurring twice. The frequencies for each occurrence category were summed across all $10^6$ simulations.

Across the $10^6$ simulations, a total of 96,982,390 distinct observed code patterns were counted. Of these, 90,283,978 (93.09 %) occurred once, 6,390,077 (6.59 %) occurred twice, 297,754 (0.307 %) occurred three times, 10,305 (0.0106 %) occurred four times, 270 (0.000278 %) occurred five times, and 6 (0.00000619 %) occurred six times (Figure S7). No patterns occurred more than six times. The simulated distribution closely agreed with the theoretical distribution predicted under equal-probability random assignment.

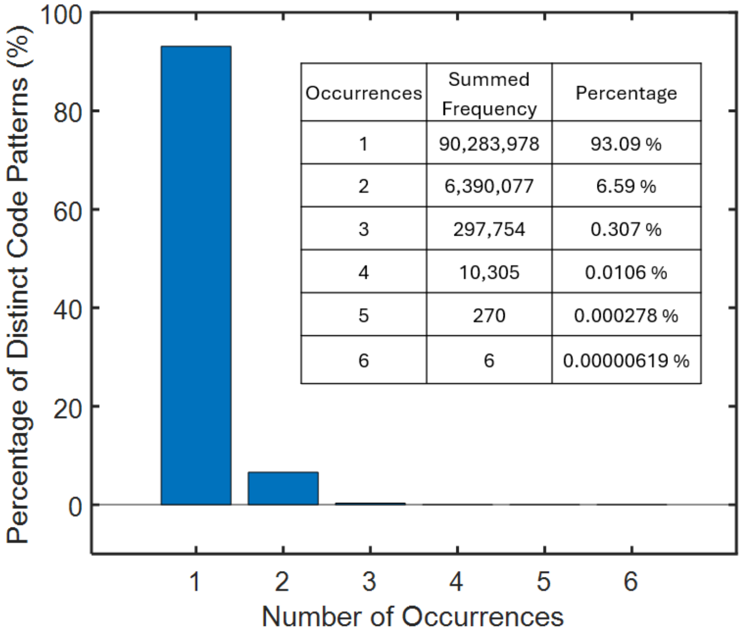


| Occurrences | Summed Frequency | Percentage |
|---|---|---|
| 1 | 90,283,978 | 93.09 % |
| 2 | 6,390,077 | 6.59 % |
| 3 | 297,754 | 0.307 % |
| 4 | 10,305 | 0.0106 % |
| 5 | 270 | 0.000278 % |
| 6 | 6 | 0.00000619 % |

**Figure S13.** Monte Carlo simulation of ternary-code pattern recurrence under random assignment. In each of $10^6$ simulations, 104 analytes were independently assigned with equal probability to the 729 possible six-digit ternary codes. Distinct code patterns occurring at least once within each simulation were categorized according to their number of occurrences. Percentages were calculated from the 96,982,390 distinct observed code patterns accumulated across all simulations. The inset table provides the corresponding summed frequencies and percentages for each occurrence category.

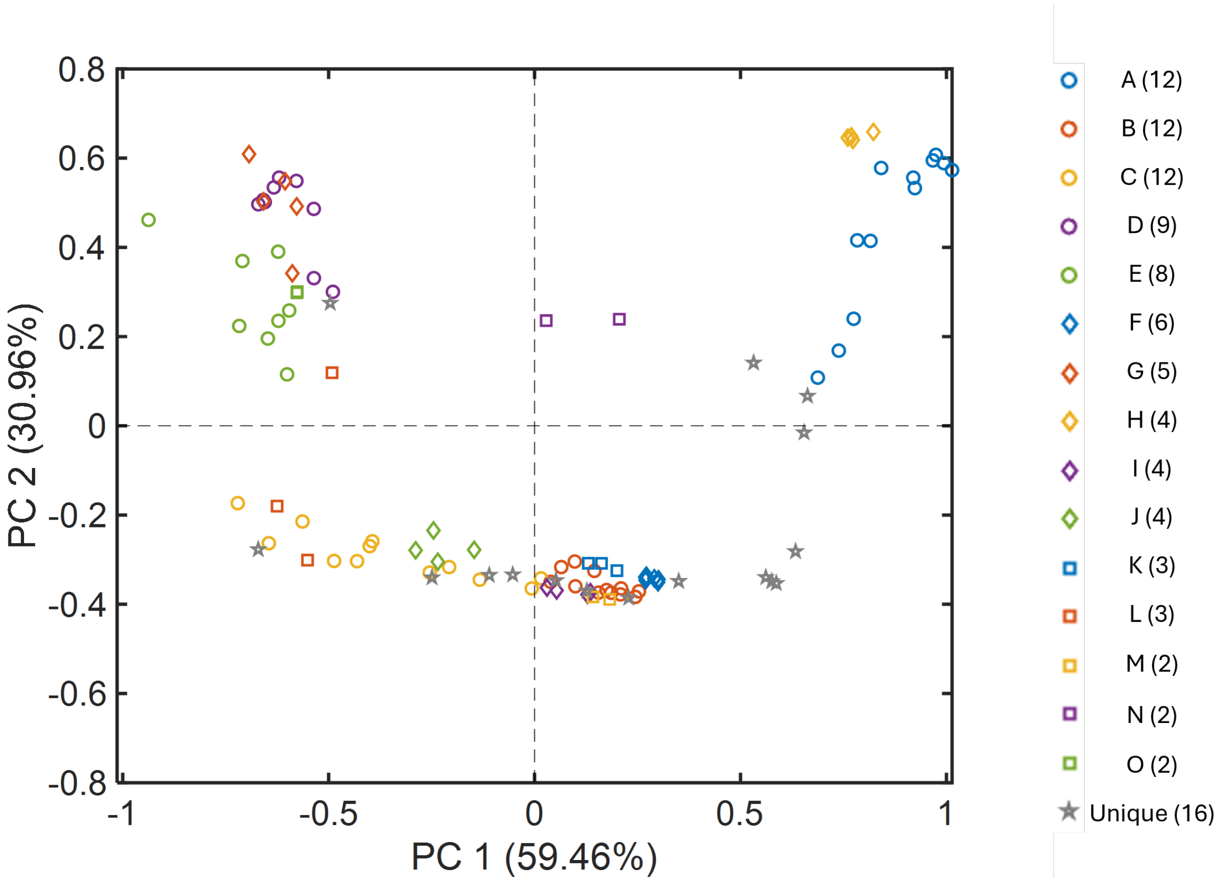


**Figure S14.** PCA of the 104 hits meeting the *p*-value < 0.01 criterion in one-way ANOVA. Tile signal intensity for each hit was normalized to the maximum value across the four sample classes (M, PA, AF, and CC), yielding values from 0 to 1. Points are colored and shaped according to ternary-code group for the 15 patterns occurring in two or more hits (A-O; n indicated in the legend). The remaining 16 hits with unique, singly occurring ternary-code patterns are shown as gray stars and labeled "Unique."

**Table S1.** Ternary-code patterns observed among the 94 hits with p-value ≥ 0.01, showing the number of occurrences for each pattern and the corresponding Monte Carlo frequency (%) under random assignment. Note: Monte Carlo frequencies were calculated from $10^6$ simulations. Patterns occurring seven or more times were not observed. (N.O., not observed)

| Number of Occurrences | Monte Carlo Frequency (%) | Ternary Pattern | | | | | |
|---|---|---|---|---|---|---|---|
| | | M vs. PA | M vs. AF | M vs. CC | PA vs. AF | PA vs. CC | AF vs. CC |
| 43 | N.O. | 0 | 0 | 0 | 0 | 0 | 0 |
| 9 | N.O. | 0 | 0 | 0 | 0 | 0 | 1 |
| 6 | $6.19\times10^{-6}$ | 0 | -1 | 0 | 0 | 0 | 0 |
| 6 | $6.19\times10^{-6}$ | 0 | 0 | 1 | 0 | 0 | 0 |
| 6 | $6.19\times10^{-6}$ | 0 | 0 | 1 | 0 | 0 | 1 |
| 6 | $6.19\times10^{-6}$ | 0 | 0 | 0 | -1 | 0 | 1 |
| 3 | 0.307 | 0 | -1 | 0 | -1 | 0 | 0 |
| 2 | 6.59 | 0 | 0 | 0 | -1 | 0 | 0 |
| 2 | 6.59 | -1 | -1 | 0 | 0 | 0 | 0 |
| 2 | 6.59 | -1 | -1 | 0 | 0 | 0 | 1 |
| 2 | 6.59 | 0 | -1 | 0 | -1 | 0 | 1 |
| 2 | 6.59 | 0 | 0 | 0 | 0 | 1 | 1 |
| 1 | 93.09 | 0 | 0 | 0 | 0 | 1 | 0 |
| 1 | 93.09 | 1 | 0 | 0 | 0 | 0 | 0 |
| 1 | 93.09 | -1 | -1 | -1 | 0 | 0 | 0 |
| 1 | 93.09 | 0 | 0 | 1 | -1 | 0 | 1 |
| 1 | 93.09 | 1 | 0 | 1 | 0 | 0 | 0 |

**Table S2.** Ternary-code patterns observed among the 104 hits with p-value < 0.01, showing the number of occurrences for each ternary-code pattern and the corresponding Monte Carlo frequency (%) under random assignment. Note: Monte Carlo frequencies were calculated from $10^6$ simulations. Patterns occurring seven or more times were not observed. (N.O., not observed)

| Number of Occurrences | Monte Carlo Frequency (%) | Ternary Pattern | | | | | |
|---|---|---|---|---|---|---|---|
| | | M vs. PA | M vs. AF | M vs. CC | PA vs. AF | PA vs. CC | AF vs. CC |
| 1 | 93.09 | 0 | 0 | 0 | -1 | 0 | 0 |
| 4 | $1.06\times10^{-2}$ | 0 | 0 | 0 | -1 | 0 | 1 |
| 3 | 0.307 | 0 | -1 | 0 | 0 | 0 | 1 |
| 1 | 93.09 | -1 | -1 | 0 | 0 | 0 | 0 |
| 1 | 93.09 | 0 | 0 | 1 | 0 | 0 | 1 |
| 1 | 93.09 | 1 | 0 | 1 | 0 | 0 | 0 |
| 12 | N.O. | 0 | -1 | 0 | -1 | 0 | 1 |
| 9 | N.O. | 1 | 1 | 1 | 0 | 0 | 0 |
| 3 | 0.307 | 1 | 0 | 1 | 0 | 0 | 1 |
| 1 | 93.09 | -1 | -1 | -1 | 0 | 0 | 0 |
| 1 | 93.09 | -1 | 0 | -1 | 1 | 0 | 0 |
| 1 | 93.09 | -1 | 0 | 0 | 1 | 1 | 0 |
| 1 | 93.09 | 1 | 0 | 0 | -1 | 0 | 1 |
| 12 | N.O. | -1 | 0 | -1 | 1 | 0 | -1 |
| 12 | N.O. | 1 | 0 | 1 | -1 | 0 | 1 |
| 6 | $6.19\times10^{-6}$ | -1 | -1 | 0 | 0 | 1 | 1 |
| 5 | $2.78\times10^{-4}$ | 1 | 1 | 1 | 0 | 1 | 0 |
| 2 | 6.59 | 0 | -1 | 1 | -1 | 0 | 1 |
| 2 | 6.59 | 0 | 1 | 1 | 1 | 1 | 0 |
| 2 | 6.59 | 1 | 1 | 1 | 0 | 0 | 1 |
| 1 | 93.09 | -1 | -1 | -1 | 0 | -1 | 0 |
| 1 | 93.09 | -1 | -1 | -1 | 0 | 0 | 1 |
| 1 | 93.09 | -1 | -1 | -1 | 0 | 1 | 0 |
| 1 | 93.09 | 1 | -1 | 0 | -1 | 0 | 1 |
| 8 | N.O. | 1 | 1 | 1 | -1 | 0 | 1 |
| 4 | $1.06\times10^{-2}$ | -1 | 0 | -1 | 1 | 1 | -1 |
| 4 | $1.06\times10^{-2}$ | 1 | -1 | 1 | -1 | 0 | 1 |
| 1 | 93.09 | -1 | -1 | -1 | 1 | 1 | 0 |
| 1 | 93.09 | 1 | 0 | 1 | -1 | 1 | 1 |
| 1 | 93.09 | 1 | 1 | 1 | 0 | 1 | 1 |
| 1 | 1 | 1 | -1 | 1 | -1 | 1 | 1 |
